# Multiplex Epigenome Editing of *MECP2* to Rescue Rett Syndrome Neurons

**DOI:** 10.1101/2022.11.30.518560

**Authors:** Junming Qian, Xiaonan Guan, Bing Xie, Chuanyun Xu, Jacqueline Niu, Xin Tang, Charles H. Li, Henry M. Colecraft, Rudolf Jaenisch, X. Shawn Liu

## Abstract

Rett syndrome (RTT) is an X-linked neurodevelopmental disorder caused by loss-of-function heterozygous mutations of Methyl CpG-binding Protein 2 (*MECP2*) on the X chromosome in girls. Reactivation of the silent wild-type *MECP2* allele from the inactive X chromosome (Xi) represents a promising therapeutic opportunity for female patients with RTT. Here, we applied a multiplex epigenome editing approach to reactivate MECP2 from Xi in RTT human embryonic stem cells (hESCs) and derived neurons. Demethylation of the *MECP2* promoter by dCas9-Tet1 with target sgRNA reactivated MECP2 from Xi in RTT hESCs without detectable off-target effects at the transcriptional level. Neurons derived from methylation-edited RTT hESCs maintained MECP2 reactivation and reversed the smaller soma size and electrophysiological abnormalities, two hallmarks of RTT. In RTT neurons, insulation of the methylation edited *MECP2* locus by dCpf1-CTCF with target CRISPR RNA enhanced MECP2 reactivation and rescued RTT-related neuronal defects, providing a proof-of-concept study for epigenome editing to treat RTT and potentially other dominant X-linked diseases.

**One-Sentence Summary:** Reactivation of *MECP2* from the inactive X chromosome by multiplex epigenome editing rescues Rett syndrome neurons in vitro.

## Introduction

Rett syndrome (RTT) is an X-linked postnatal progressive neurodevelopmental disorder associated with severe mental disability and autism-like syndromes that manifests in girls during early childhood (*1, 2*). RTT is caused by loss-of-function mutations of the Methyl CpG-binding Protein 2 gene (*MECP2*) on the X chromosome (*3*). Most female patients with RTT carry a heterozygous mutation of *MECP2* in which a wild-type (WT) allele and a loss-of-function allele are randomly inactivated during development, resulting in ∼50% of neurons in the patient without functional MeCP2 protein (*1, 4, 5*). Mice carrying null alleles of *Mecp2* closely mimic symptoms seen in patients, including microcephaly, hindlimb clasping corresponding to repetitive hand movements seen in patients, irregular breathing with apneas and shortened lifespan, and are faithful models of the disease (*6-10*). The development of RTT-like symptoms in mice can be halted or even reversed in the adult after genetic or viral restoration of MeCP2 protein expression (*11-15*). Thus, reactivation of the silenced WT allele of *MECP2* from the inactive X chromosome (Xi) represents an exciting research direction with promising therapeutic opportunity for RTT (*16, 17*), as it attacks the root cause of this potentially reversible disease by restoring MeCP2 expression. However, attempts to restore MeCP2 with a single non-targeted approach such as treatment with DNA methylation inhibitors or anti-sense oligo (ASO) targeting the long non-coding RNA XIST required for X chromosome inactivation only resulted in limited MECP2 reactivation with potential off-target toxicity (*16*).

Given X chromosome inactivation is a classic biological process mediated by a cascade of epigenetic events including decoration by long non-coding RNA *XIST*, DNA methylation, histone modifications, and reorganization of the 3D chromosomal structure by architectural proteins such as CTCF (*18*), we hypothesized that precise manipulation of the epigenetic status of the silenced *MECP2* allele on Xi could reactivate its expression. We and other laboratories developed a series of epigenome editing tools that allows for precisely manipulating the epigenetic status of targeted genomic loci (*17, 19, 20*). These editing tools consist of a DNA targeting module such as a catalytically dead CRISPR/Cas protein (*21, 22*) and an epigenetic modifier module such as DNA methylation/demethylation enzymes Dnmt or Tet (*23*). Such fusion proteins can be targeted to the specific loci by single guide RNA (sgRNA) to mediate epigenetic modifications (*17, 19*). For instance, we applied dCas9-Tet1 to specifically demethylate the hypermethylated CGG repeat expansion mutation in the 5’UTR region of *FMR1* gene that causes Fragile X syndrome (FXS), demonstrating that this targeted demethylation reactivated FMR1 and rescued the neuronal defects of FXS neurons (*24*). Here, to overcome the multiple layers of epigenetic silencing on Xi, we applied a DNA methylation editing tool (dCas9-Tet1) and developed a DNA insulating tool (dCpf1-CTCF) to reactivate the *MECP2* allele from Xi and rescue the pathological phenotype of RTT neurons.

## Result

### Reactivation of a MECP2 reporter on Xi by DNA methylation editing

To distinguish the *MECP2* alleles on active X chromosome (Xa) and inactive X chromosome (Xi) during reactivation experiments, we used a MECP2 dual-color reporter human embryonic stem cell (hESC) line (29-R) that was genetically engineered from a wild-type female hESC line (NIH registration number: WIBR-2, #29) (*25*). This 29-R hESC line has green fluorescent protein (GFP) inserted in-frame after Exon 3 at the *MECP2* locus on the Xi and tdTomato similarly inserted at the *MECP2* locus on the Xa, and thus only expresses tdTomato but not GFP (**Figure 1A)**. We also included a 29-G hESC line (*25*) with the GFP reporter on Xa and tdTomato on Xi as a positive control to evaluate the reactivation efficacy of the MECP2 GFP reporter in edited 29-R cells. Because the polyA signal sequence was engineered after GFP and tdTomato, both 29-R and 29-G cells do not express MeCP2 proteins and thus reactivation of the MECP2 reporter from Xi would not influence the downstream target genes of MeCP2. To identify the target region at the *MECP2* locus for DNA methylation editing, we compared the DNA methylation status of female (Xa;Xi) and male (Xa) hESC cells in a published database (*26*). We identified a 1.7 kb differentially methylated region (DMR) in the *MECP2* promoter region that is methylated in female cells (about 50% in average due to one methylated allele on Xi and one unmethylated allele on Xa) but not in male cells (due to only one unmethylated allele on Xa; **Figure 1B**). We reasoned that targeted demethylation of this region would activate the *MECP2* allele from Xi (Figure 1A). Thus, based on the effective range for methylation editing (*20*), we designed ten sgRNAs to target dCas9-Tet1 to this differentially methylated region, and four Pyro-seq assays to examine the methylation of four areas (a, b, c, and d) within this region. We delivered dCas9-Tet1-P2A-BFP (dC-T) and ten target sgRNAs with mCherry reporter via lentiviral transduction into 29-R hESCs, and FACS-isolated infection-positive cells (BFP+;mCherry+) for downstream analysis. The DNA methylation percentages of MECP2 promoter in #29 (the parental line for 29-R and 29-G), mock 29-R, and mock 29-G cells were around 30-50%, and reduced to ∼7% in 29-R cells expressing dC-T and target sgRNAs for all the four Pyro-seq areas within the targeted differentially methylated region (**Figure 1C)**, suggesting efficient demethylation was achieved in the targeted cells. In contrast, the DNA methylation percentage of this differentially methylated region in 29-R cells expressing dCas9 fused with a catalytically dead Tet1 (dC-dT) with the same group of target sgRNAs did not change, confirming the decreased DNA methylation was due to the targeted dC-T enzymatic activity. As a consequence of targeted demethylation of the *MECP2* promoter, the edited 29-R cells (labeled as 29-R_dC-T+10), but not the control 29-R cells (labeled as 29-R_dC-dT+10), turned on the GFP reporter for MECP2 on Xi (**Figure 1D**), suggesting demethylation of its promoter can reactivate MECP2 from Xi in hESCs. To examine the reactivation efficacy for each individual sgRNA, we generated a doxycycline (Dox)-inducible dCas9-Tet1 expression 29-R cell line (labeled as 29-R_138-dC-T) validated by qPCR analysis (**Figure 1E)**. We infected this cell line with each lentiviral sgRNA (labeled as 1, 2, 3, 4, 5, 6, 7, 8, 9, 10 individually) or a mixture of ten sgRNAs (labeled as 1-10), and similar expression quantities of sgRNA-mCherry as well as dCas9-Tet1 upon Dox treatment among these samples were validated by qPCR (**Figure S1**). Pyro-seq of these cells showed sgRNA-3 resulted in the most robust demethylation (18% as the average DNA methylation percentage of each CpG within the targeted *MECP2* promoter) compared to the average of 44% in unedited 29-R cells (**Figure 1F**). Reporter expression analysis of these cells by qPCR showed sgRNA-3 resulted in a similar amount of GFP reactivation compared to the effect by the mixture of ten sgRNAs (**Figure 1G**), suggesting that sgRNA-3 is sufficient to completely reactivate the *MECP2* reporter from Xi at the hESC stage. For the following experiments, we used sgRNA-3 for off-target effect analysis and DNA methylation editing.

**Figure 1.**
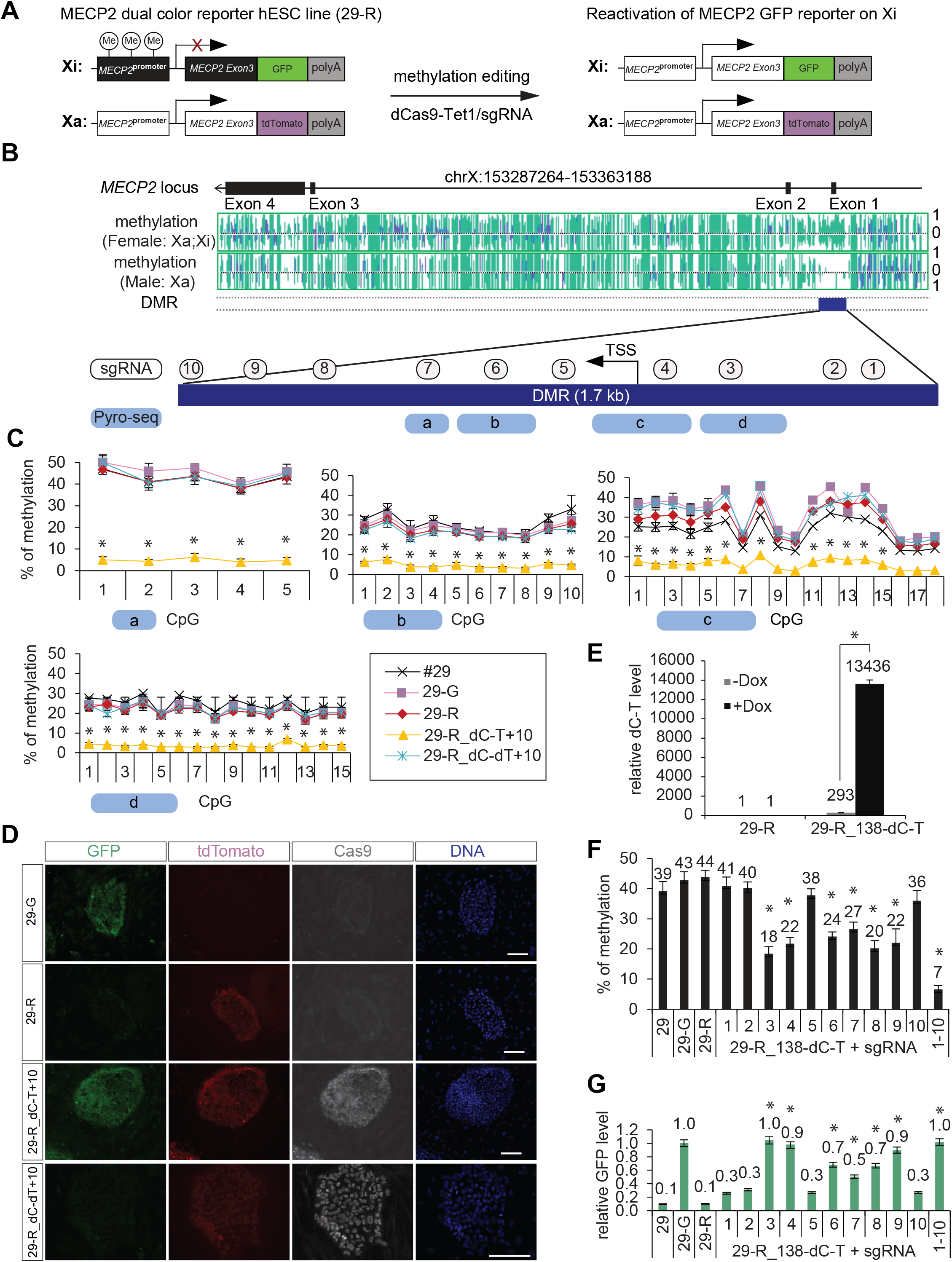
Reactivation of the MECP2 reporter on Xi by DNA methylation editing. (**A**) Scheme of genetically engineered MECP2 dual color reporter hESC lines derived from a wild-type female hESC (NIH registration code: WIBR-1, #29) after methylation editing. For the 29-R cell line, GFP was inserted after MECP2 exon 3 in frame followed by poly-A termination signal on the inactive X chromosome (Xi), and tdTomato was inserted after MECP2 exon 3 in frame followed by poly-A termination signal on the active X chromosome (Xa). For 29-G cell line, GFP is on Xa and tdTomato is on Xi. (**B**) Illustration of the differentially methylated region (DMR) between female and male hESCs at the *MECP2* promoter. Ten sgRNA were designed to target this DMR, and four pyro-sequencing (Pyro-seq) assays were designed to measure the DNA methylation of this DMR. (**C**) 29-R hESCs were infected with lentiviruses expressing dCas9-Tet1-P2A-BFP (dC-T) or dCas9-dead Tet1-P2A-BFP (dC-dT) with ten sgRNAs with mCherry as a fluorescent marker targeting this DMR at the *MECP2* promoter as illustrated in (B). The infection positive cells (BFP+;mCherry+) were isolated by FACS and subject to Pyro-seq analysis. Shown is the mean percentage ± SD of three biological replicates. (**D**) Immunofluorescence (IF) staining of cells described in (C) with antibodies against GFP and Cas9. Scale bar: 300 um. (**E**) A Dox-inducible dCas9-Tet1 expression cassette was inserted into the 29-R hESCs via PiggyBac transposon system (labeled as 29-R_138-dC-T), and dCas9-Tet1 expression was examined by RT-qPCR in response to Dox treatment. Shown is the mean ± SD of three biological replicates. (**F**) Cells in (E) were infected with individual target sgRNA (labeled 1 to 10) or the mixture of ten sgRNA (labeled 1-10) targeting the *MECP2* promoter and then subjected to methylation analysis by Pyro-seq in the presence of Dox. Shown is the average DNA methylation of CpGs within the targeted *MECP2* promoter region ± SD of three biological replicates. (**G**) RT-qPCR analysis of GFP reporter for MECP2 on Xi in cells in (F). GFP expression was normalized to 29-G cells with GFP reporter for MECP2 on Xa. Shown is the mean percentage ± SD of at least two biological replicates. \**P* < 0.05, one-way analysis of variance (ANOVA) with Bonferroni correction.

### Methylation editing efficacy and off-target effects

Potential off-target effect is a critical parameter that needs to be evaluated for epigenome editing approach. Towards this goal, we performed a genome-wide analysis at the DNA methylation and transcriptional levels for DNA methylation editing of the *MECP2* promoter by dCas9-Tet1 with sgRNA-3. Although we identified 27 genome-wide binding sites for dCas9-Tet1/sgRNA-3 by anti-Cas9 chromatin immunoprecipitation sequencing (ChIP-seq) using edited 29-R hESCs (**Figure 2A, Data file S1**), the targeted MECP2 promoter showed the highest enrichment of dCas9-Tet1 binding, suggesting a high targeting specificity enabled by this sgRNA. Then, we performed anti-Cas9 ChIP-Bisulfite-seq to compare the DNA methylation percentages of these 27 binding sites between 29-R cells expressing dCas9-Tet1/sgRNA-3 (methylation-edited sample) and dCas9-dTet1/sgRNA-3 (control sample). As shown in **Figure 2B** and **Data file S2**, the targeted MECP2 promoter showed the largest change of DNA methylation (51% reduction in edited 29-R cells), demonstrating a high efficacy of targeted methylation editing. Among the remaining 26 sites, 21 binding sites labeled by empty circles showed a change of DNA methylation < 16%, and 5 other binding sites labeled by red color circles showed a change of DNA methylation between 16% to 51%. Notably, the RNA-seq result in **Figure 2C** and **Data files S3-S5** showed robust reactivation of the GFP reporter for MECP2 on Xi (17-fold) in methylation-edited 29-R cells, but no significant changes (adjusted *P* value < 0.01 and fold change > 2) in the expression of genes associated with the other 26 binding sites by dCas9-Tet1/sgRNA-3, suggesting no detectable off-target effect at the transcriptional level. To further evaluate a potential off-target effect of dCas9-Tet1 on genome-wide methylation, we performed whole-genome bisulfite sequencing. Our protocol included three biological replicates for each of the experimental groups in which 29-R cells express dCas9-Tet1 (dC-T) or dCas9-dead Tet1 (dC-dT) with sgRNA-3, and covered 20,391,442 CpG sites (at least 5x reads coverage per CpG) representing 72% of the total 28,299,634 CpG sites in the human genome. The average CpG methylation in the dC-T group and dC-dT group were the same (76%; **Figure S2A**), suggesting no alteration on the overall DNA methylation by dC-T. Among these sequenced 20,391,442 CpG sites (**Figure S2B)**, 99.96% of CpGs showed no significant change in DNA methylation (adjusted *P* value < 0.01 and change in methylation > 20%), and only 0.04% of these CpGs (7740 cytosines) including the *MECP2* locus showed changes in DNA methylation larger than 20%, termed differentially methylated CpGs (**Figure S2C**). Among these 7740 differentially methylated CpGs, 2307 cytosines that showed higher methylation percentages in dC-T compared to dC-dT samples were not considered off-target effects by dC-T, but instead likely caused by differential epigenetic drift during cell passaging of hESCs (*27*). We mapped the remaining 5433 CpGs that showed less methylation in dC-T compared to dC-dT samples onto 239 genes that contain at least three differentially methylated CpGs with the distance between each differentially methylated CpG less than 250 bp. Among these 239 genes (**Data file S6**), 220 of them did not show changes in gene expression, whereas 19 of them showed significant changes of expression (adjusted *P* value < 0.01 and fold change > 2). However, none of these 19 genes were bound by dCas9-Tet1 as examined by ChIP-seq (**Figure 2C**), suggesting the expression changes of these 19 genes were unlikely caused by dCas9-Tet1. In summary, analyses of whole-genome bisulfite sequencing, ChIP-seq, and RNA-seq results did not detect off-target effects by dCas9-Tet at the transcriptional level.

**Figure 2.**
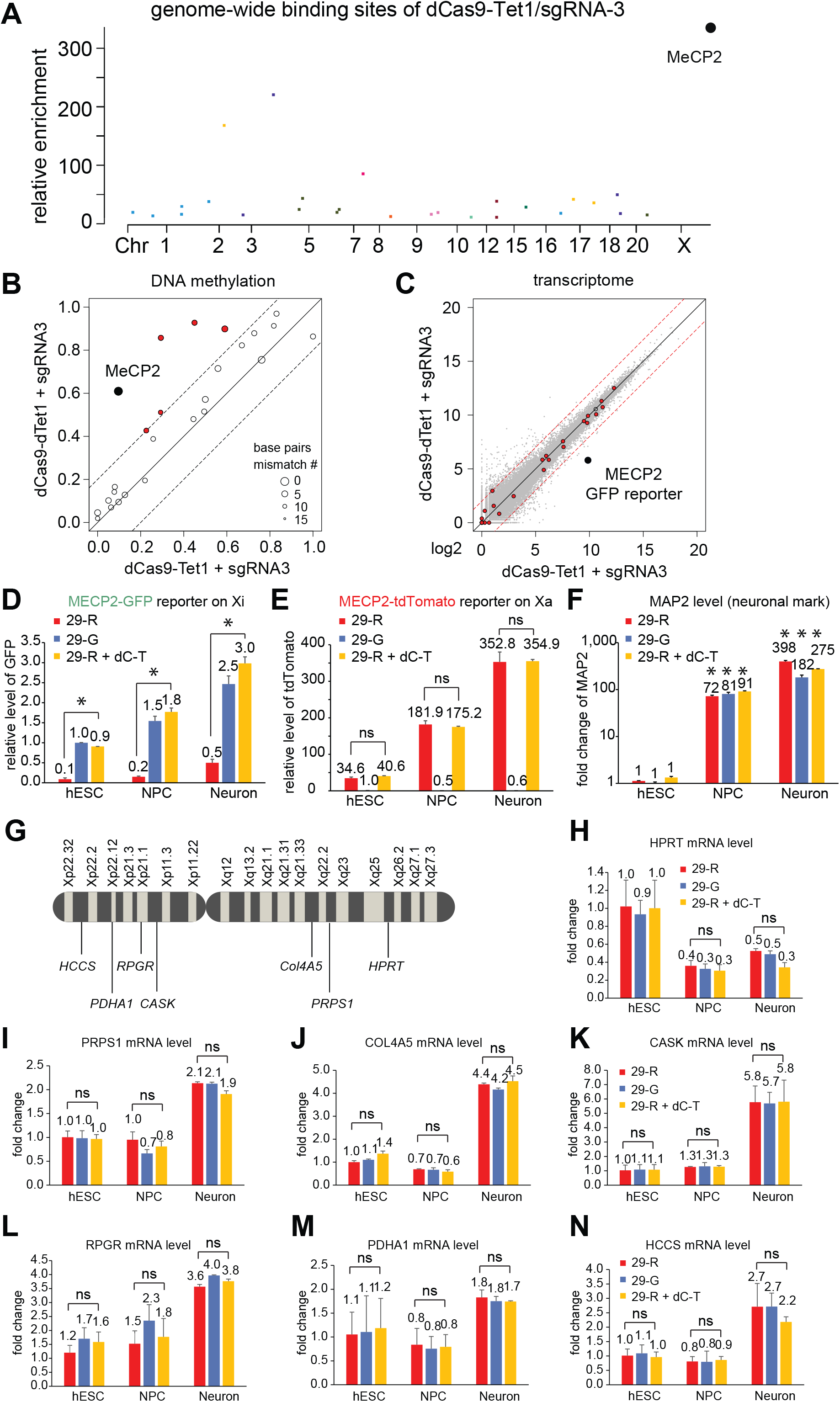
DNA methylation editing efficacy and off-target effects. (**A**) A Manhattan plot showing 27 genome-wide binding sites of dCas9-Tet1 with the *MECP2* target sgRNA-3 in 29-R cells identified by anti-Cas9 ChIP-seq. (**B**) DNA methylation of these 27 binding sites measured by anti-Cas9 ChIP-Bisulfite-seq of 29-R cells expressing dCas9-Tet1 with sgRNA-3 or dCas9-dTet1 with sgRNA-3. MECP2 is labeled in black; five binding sites with a change of methylation larger than 16% are labeled in red. The diameter of a circle is in proportion to the number of matched base pairs to the sgRNA-3 target sites. Blue lines of circles indicate binding sites overlapping with a promoter region. The dashed lines mark the 16% methylation difference between samples. (**C**) Transcriptomes of cells in (B) by RNA-seq. Red dots highlight the genes associated with the 26 dCas9-Tet1 binding sites identified in (A). MECP2-GFP reporter on Xi is labeled with a black dot. Dashed red lines mark the 2-fold difference between the samples. (**D-F**) Mock 29-R, mock 29-G, or methylation-edited 29-R hESCs were differentiated into neuronal precursor cells (NPCs) and neurons for gene expression analysis by qPCR. Shown is the mean ± SD of three biological replicates. (**G**) Illustration showing the location of genes on the X chromosome prone to erosion of X chromosome inactivation in high passage of female hESC/iPSC. (**H-N**) Gene expression analysis of cells in (D) by qPCR: *HPRT* in **H**, *PRPS1* in **I**, *COL4A5* in **J**, *CASK* in **K**, *RPGR* in **J**, *PDHA1* in **M**, and *HCCS* in **N**. Shown is the mean ± SD of two biological replicates. \**P* < 0.05, one-way ANOVA with Bonferroni correction. The differences between 29-R and 29-R_dC-T+sgRNA in **H**-**N** were not significant (ns; *P* > 0.05).

Erosion of X chromosome inactivation was observed in high-passage female hESCs and induced pluripotent stem cells (iPSCs), with derepression of X-linked genes on Xi including *HPRT, RPGR, PRPS, PDHA1, COL4A5, HCCS*, and *CASK* (*28*). To test whether the erosion of X chromosome inactivation occurred in the methylation-edited 29-R hESCs, potentially confounding our conclusion on MECP2 reactivation by methylation editing, we examined the expression of the GFP and tdTomato reporters on Xi and Xa for MECP2 and these seven X-linked genes in methylation-edited 29-R hESCs, neuronal precursor cells (NPCs), and cortical neurons derived from edited 29-R hESCs using a well-established neuronal differentiation protocol (*29*). The GFP reporter for *MECP2* on Xi remained active in NPCs and neurons (**Figure 2D)**, suggesting that MECP2 reactivation can be maintained during neuronal differentiation. The tdTomato reporter for the Xa *MECP2* allele was expressed to a similar extent as in edited 29-R cells compared to mock 29-R cells during neuronal differentiation (**Figure 2E)**, suggesting that methylation editing did not affect the expression of the active *MECP2* allele on Xa. The gradual increase of neuronal mark MAP2 from hESCs to NPCs and neurons confirmed robust neuronal cellular differentiation (**Figure 2F**). All seven X-linked genes (**Figure 2G**) prone to the erosion of X chromosome inactivation showed no difference in expression between mock 29-R, mock 29-G, and methylation-edited 29-R hESCs, or in NPCs and neurons derived from these hESCs (**Figure 2H-N**). These results demonstrate that reactivation of the GFP reporter for MECP2 on Xi is not the consequence of X chromosome inactivation erosion in methylation-edited 29-R cells.

### Functional rescue of RTT neurons derived from edited RTT-like hESCs

To evaluate the functional consequences of MECP2 reactivation by DNA methylation editing, we used an RTT-like hESC (#860) line (*25*) that is genetically engineered from a wild-type female hESC line (NIH registration code: WIBR-3). This RTT-like hESC line has a wild-type allele of *MECP2* silenced on Xi and a knock-out allele of MECP2 by a GFP-polyA stop cassette after Exon 3 on Xa and thus does not express MeCP2 protein (**Figure 3A)**. We delivered dCas9-Tet1-P2A-BFP and sgRNA-mCherry via lentiviral transduction of this RTT hESC line and isolated infection-positive (BFP+;mCherry+) cells by FACS. Gene expression analysis by qPCR targeting protein-coding Exon 4 showed a complete reactivation of the wild-type allele of *MECP2* from Xi compared to the WIBR-3 sample (labeled WT) by either the target sgRNA-3 alone or the mixture of ten sgRNAs (**Figure 3B**), consistent with our result using the MECP2 dual-color reporter 29-R hESCs. When the methylation-edited RTT hESCs were differentiated into neurons, Western blot experiments showed the restoration of MeCP2 protein in the neurons with dC-T and sgRNA-3 to 82% of the MECP2 protein abundance in neurons derived from the parental line of WIBR-3 hESCs with wild-type MECP2 alleles on Xa and Xi (**Figure 3C**).

**Figure 3.**
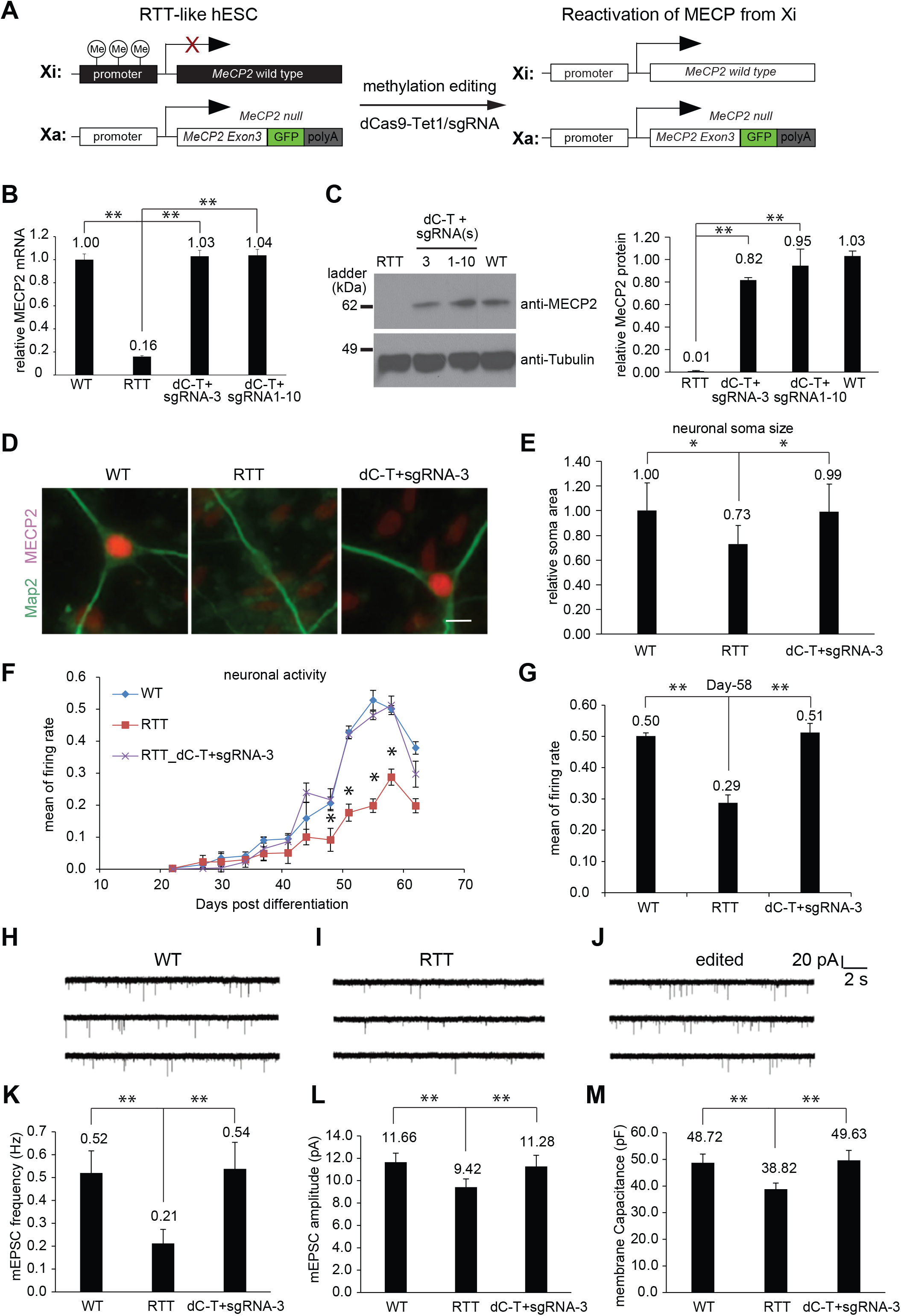
Functional rescue of RTT neurons derived from edited RTT hESCs. (**A**) Scheme of an RTT-like hESC line (#860) genetically engineered from a wild-type female human ESCs (NIH registration code: WIBR-3) after methylation editing. In this RTT-like cell line, the wild-type allele of *MECP2* is on Xi, and the *MECP2* null function allele is on Xa. (**B**) RTT-like hESCs (labeled as RTT) were infected with lentiviral dCas9-Tet1-P2A-BFP with either sgRNA-3 alone or 10 sgRNAs together. Infection-positive cells were isolated by FACS and subjected to RT-qPCR analysis of *MECP2* expression with primers targeting the exon 4 region that will only be expressed from the wild-type allele on Xi but not the null function allele on Xa. The expression of *MECP2* mRNA in these samples was normalized to *WIBR-3* (labeled WT). Shown is the mean ± SD of three biological replicates. (**C**) Western blot analysis of the neurons derived from the cells described in (B). Protein abundance of MeCP2 was quantified by ImageJ and is shown as the mean of relative percentages as compared to WT neurons ± SD of two biological replicates. (**D**) Neurons in (C) were grown on mouse astrocytes to promote neuronal maturation and then IF stained with anti-MeCP2 and anti-Map2 antibodies. Scale bar: 30 um. (**E**) Soma sizes of neurons in (D) were quantified by ImageJ. (**F**) Neurons in (C) were grown on the multi-electrode array plate for measurement of electrophysiological activities along the neuronal maturation process. Shown is the mean ± SD of biological replicates for each group of neurons. (**G**) Neuronal activities of neurons in (F) on the day of maturation (day 58). (**H-J**) Representative trace images showing spontaneous synaptic events of neurons in (D). (**K to M**) The mEPSC frequency (**K**), mEPSC amplitude (**L**), and membrane capacitance (**M**) of neurons in (D). Shown is the mean ± SD of at least two biological repeats with more than 20 neurons for each condition. \**P* < 0.05, \*\**P* < 0.01, one-way ANOVA with Bonferroni correction.

Then, we examined whether the restoration of MeCP2 protein in RTT neurons rescued neuronal defects including smaller soma size and abnormal electrophysiological properties. To quantify neuronal soma size, we differentiated wild-type control WIBR-3 hESCs, mock RTT hESCs, and methylation-edited RTT hESCs by dC-T and sgRNA-3 into cortical neurons using a well-established differentiation protocol (*30*). We cultured these neurons on mouse astrocytes for eight weeks to promote neuronal maturation (*31*), and then performed immunofluorescence (IF) staining with antibodies against MeCP2 and a neuronal marker MAP2 to outline the soma and neuronal processes. MeCP2 protein was detected in the neurons derived from WT and edited RTT hESCs but not mock RTT hESCs (**Figure 3D)**. Quantification of soma sizes by Image J with more than 100 neurons from each group showed that the smaller soma size defected in RTT neurons (73% of WT) was rescued in neurons derived from edited RTT hESCs (99% of WT) (**Figure 3E)**. To examine spontaneous electrophysiological activity during neuronal maturation, we performed a multi-electrode array time course assay to measure the firing rates of these neurons. RTT neurons displayed lower firing rates than WT neurons along the neuronal maturation process, but neurons from edited RTT hESCs showed indistinguishable firing rates compared to WT neurons (**Figure 3F)**, suggesting restoration of spontaneous electrophysiological activity in these RTT neurons by methylation editing. Quantification of the firing rates on the day of neuronal maturation (peak of firing rate on Day 58 post-differentiation) confirmed the successful rescue of the neurons from edited RTT hESCs (**Figure 3G**). To evaluate the rescue effect on RTT electrophysiological defects at single-neuron resolution, we cultured these neurons on mouse astrocytes for eight weeks to ensure functional maturation, and then performed patch-clamp recording to examine electrophysiological properties including the frequency and amplitude of mini excitatory postsynaptic current (mEPSC) and membrane capacitance. Representative recording trace images (**Figure 3H-J)** and quantified results (**Figure K-M)** show that neurons differentiated from edited RTT hESCs displayed a similar mEPSC frequency (0.54 Hz) and amplitude (11.28 pA) compared to the 0.52 Hz and 11.66 pA in WT neurons, indicating the rescue of the number and strength of excitatory synapses in these neurons. The membrane capacitance in the neurons from edited RTT hESCs (49.63 pF) was also restored in comparison to the 48.72 pF in WT and 38.82 in RTT neurons (**Figure 3M**), consistent with our soma size measurements. In summary, we conclude that functional rescue of RTT-associated neuronal defects occurred in neurons derived from DNA methylation edited RTT-like hESCs.

### Direct reactivation of MECP2 in RTT neurons

We sought to directly reactivate the *MECP2* allele from Xi in RTT neurons by DNA methylation editing, which if successful would expand the potential therapeutic window for the treatment of RTT. We delivered dCas9-Tet1-P2A-BFP and target sgRNA-3 via lentiviral transduction into neurons derived from 29-R MECP2 reporter hESCs, and performed qPCR analysis of infected 29-R neurons to quantify the reactivation of GFP reporter for MECP2 on Xi in a time-course experiment. We detected 50% reactivation of GFP reporter for MECP2 on Xi after 10 days post-infection, and then the reactivation decreased to ∼30% after 15 days and maintained at around 30% on day 20 (**Figure 4A)**, suggesting that DNA methylation editing of the *MECP2* promoter by dCas9-Tet1 can partially reactivate the *MECP2* allele from Xi in RTT neurons.

**Figure 4.**
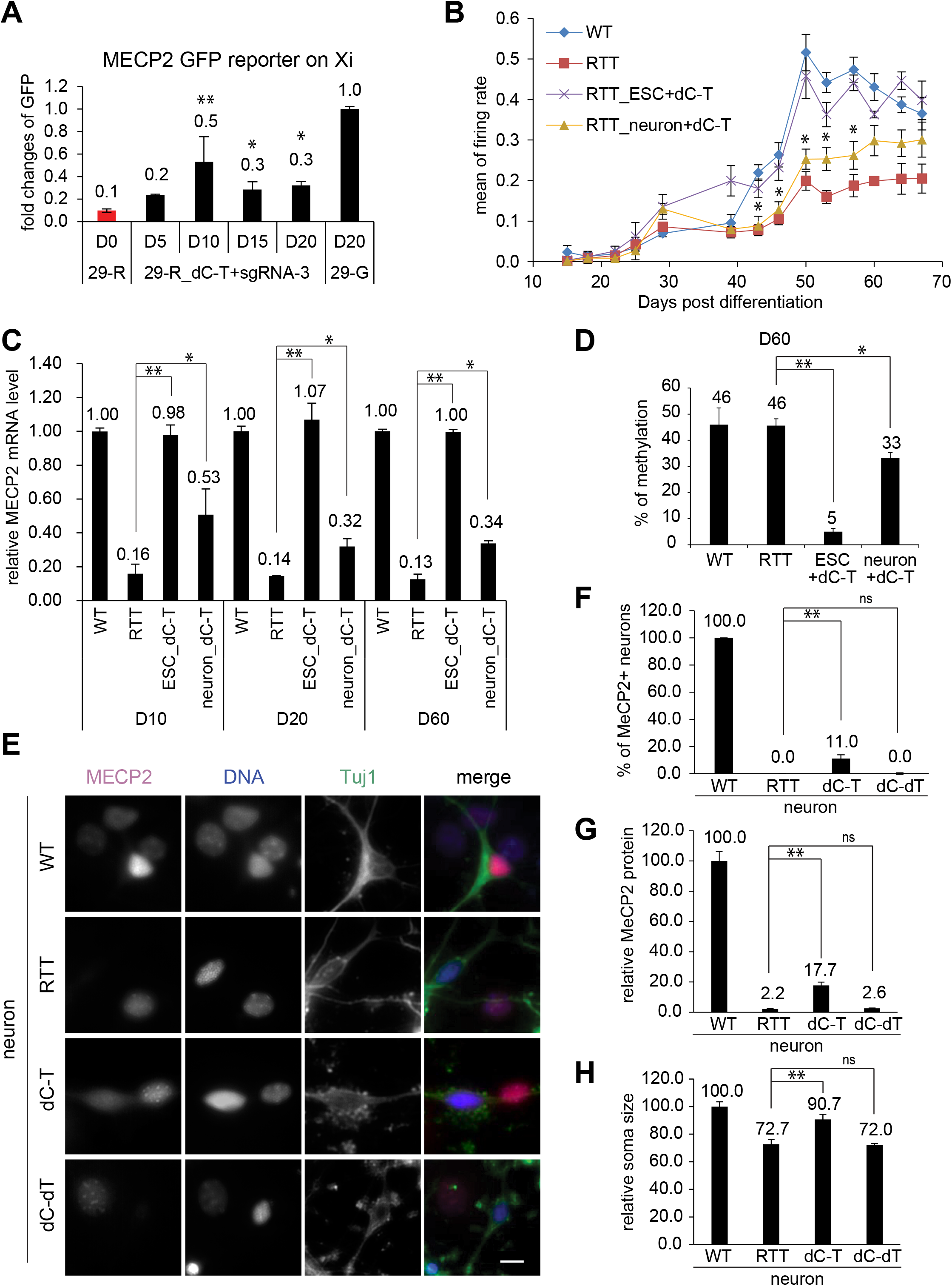
Direct editing and reactivation of MECP2 in RTT neurons. (**A**) Neurons derived from 29-R hESCs were infected with lentiviral dCas9-Tet1 (dC-T) and sgRNA-3, and then subjected to qPCR analysis on Day 5 (D5), Day 10 (D10), Day 15 (D15), and Day 20 (D20) after infection. GFP expression was normalized to 29-G neurons with GFP reporter for MECP2 on Xa. Shown is the mean ± SD of biological triplicates for each time point. (**B**) Neurons derived from wild-type WIBR-3 hESCs (labeled WT), mock RTT-like hESCs (labeled RTT), RTT-like hESCs edited by dCas9-Tet1/sgRNA-3 (labeled RTT_ESC+d-T), or RTT neurons infected by lentiviral dCas9-Tet1 and sgRNA-3 (labeled RTT_neuron+dC-T) were grown on the MEA plate for measurement of firing rates along the neuronal maturation process. Shown is the mean ± SD of biological triplicates for each condition. (**C**) *MECP2* mRNA expression of the neurons in (B) measured by qPCR. (**D**) DNA methylation of the *MECP2* promoter in neurons in (B) was measured by Pyro-seq on Day 60 (D60). Shown is the average DNA methylation of CpGs within the targeted *MECP2* promoter region ± SD of three biological replicates. (**E**) Neurons derived from RTT-like hESCs were infected with lentiviral dCas9-Tet1 and sgRNA-3 (labeled dC-T) or dCas9-deadTet1 and sgRNA-3 (labeled dC-dT) were grown on mouse astrocytes to promote neuronal maturation and then IF stained with anti-MeCP2 and anti-Tuj1 antibodies on Day 60. Scale bar: 30 um. (**F**) Percentages of neurons (Tuj1+) expressing MeCP2 protein within each group of samples described in (E). (**G**) Quantification of MeCP2 protein abundance in the MeCP2-positive neurons in (E). (**H**) Quantification of soma sizes of the neurons in (E). Quantifications were done using ImageJ and are shown as the mean of relative percentages as compared to WT neurons, ± SD of at least two biological replicates. \**P* < 0.05, \*\**P* < 0.01, one-way ANOVA with Bonferroni correction.

To examine whether this partial reactivation of MECP2 results in a functional rescue of RTT-associated neuronal defects, we performed a series of characterization experiments using neurons derived from RTT-like hESCsas control. First, we did a multi-electrode array to trace spontaneous electrophysiological activities during neuronal maturation. Whereas neurons derived from edited RTT hESCs (labeled RTT_ESC+dC-T) again displayed similar neuronal firing rates compared to WT neurons, RTT neurons transduced with lentiviral dCas9-Tet1 and target sgRNA-3 (labeled RTT_neuron+dC-T) only displayed a partial rescue of the lower firing rates observed in mock RTT neurons (**Figure 4B)**. Consistent with this result, gene expression analysis of these neurons at different time points along the neuronal maturation by qPCR showed 53% of reactivation of MECP2 mRNA on Day-10 post-infection and maintained as ∼30% reactivation of MECP2 from Day 20 to Day 60 (**Figure 4C)**. The lower reactivation efficacy of MECP2 from Xi is likely due to the moderate demethylation achieved by dCas9-Tet1 in these neurons (33% as the average methylation percentage of CpGs within the targeted *MECP2* promoter region) compared to 5% in neurons derived from edited RTT hESCs as measured by Pyro-seq (**Figure 4D)**. To quantify the restoration of MeCP2 protein and soma size, we cultured these neurons on mouse astrocytes to promote neuronal maturation and then IF stained with MeCP2 and neuronal marker Tuj1 antibodies. As shown by the representative images in **Figure 4E** and quantification in **Figure 4F**, IF staining of these neurons detected moderate expression of MeCP2 protein in ∼11% of RTT neurons after infection with lentiviral dC-T and sgRNA-3, but not in the mock RTT neurons or RTT neurons infected with lentiviral dC-dT and sgRNA-3. To quantify MeCP2 protein in these neurons, we used mouse astrocytes (Tuj1 negative cells) that express low amounts of MeCP2 protein within each group of samples as an internal control. Quantification using ImageJ showed that 17.7% of MeCP2 protein was restored in the MeCP2-positive RTT neurons (**Figure 4G**), and 90.7% rescue of soma size in these neurons (**Figure 4H**). These results suggest that DNA methylation editing of RTT neurons by dCas9-Tet1 can partially reactivate MECP2, resulting in a functional rescue of electrophysiological defects and the small soma size of edited RTT neurons.

### Reactivation of MECP2 and rescue of RTT neurons by multiplex epigenome editing

Next, we aimed to improve the reactivation efficacy of MECP2 from Xi in RTT neurons. Considering X chromosome inactivation is mediated by multiple epigenetic events including decoration by long non-coding RNA XIST, DNA methylation, histone modifications, and compaction of 3D chromatin structure (*18*), we hypothesized that an additional epigenetic modality can be combined with DNA methylation editing by dCas9-Tet1 to achieve a high amount of stable MECP2 reactivation from Xi in RTT neurons. We reasoned that such epigenetic changes required for MECP2 reactivation in RTT neurons could be identified by comparing mock RTT neurons with functionally rescued RTT neurons derived from methylation-edited RTT hESCs. Towards this goal we performed ChIP-seq with an antibody against the chromatin structural protein CTCF and a circular chromosome conformation capture sequencing assay (4C-seq) with a viewpoint at the *MECP2* promoter regions in these cell types. We identified two CTCF anchor sites flanking the *MECP2* locus at which the enrichment of CTCF binding were doubled in the functionally rescued RTT neurons derived from methylation edited RTT hESCs compared to mock RTT neurons (**Figure 5A)**, suggesting MECP2 reactivation-specific CTCF recruitment at these two anchor sites. This is consistent with previous studies showing the depletion of CTCF proteins from the Xi except for the maintenance of CTCF binding at the transcriptionally active escapee gene loci on Xi (*32, 33*). These two anchor sites also showed an increased enrichment of CTCF occupancy in the naïve stage of RTT hESCs with two Xa compared to that in the primed stage of RTT hESCs with one Xa and one Xi (**Figure 5A**), confirming an X chromosome activation-specific signature on CTCF recruitment. Furthermore, 4C-seq demonstrated that the genomic interaction frequency of the *MECP2* promoter decayed beyond these two CTCF anchor sites, indicating an insulation function mediated by CTCF. Considering all these observations together, the increased enrichment of CTCF binding at these two anchor sites likely contributes to the stable reactivation of MECP2 on Xi in the functionally rescued RTT neurons.

**Figure 5.**
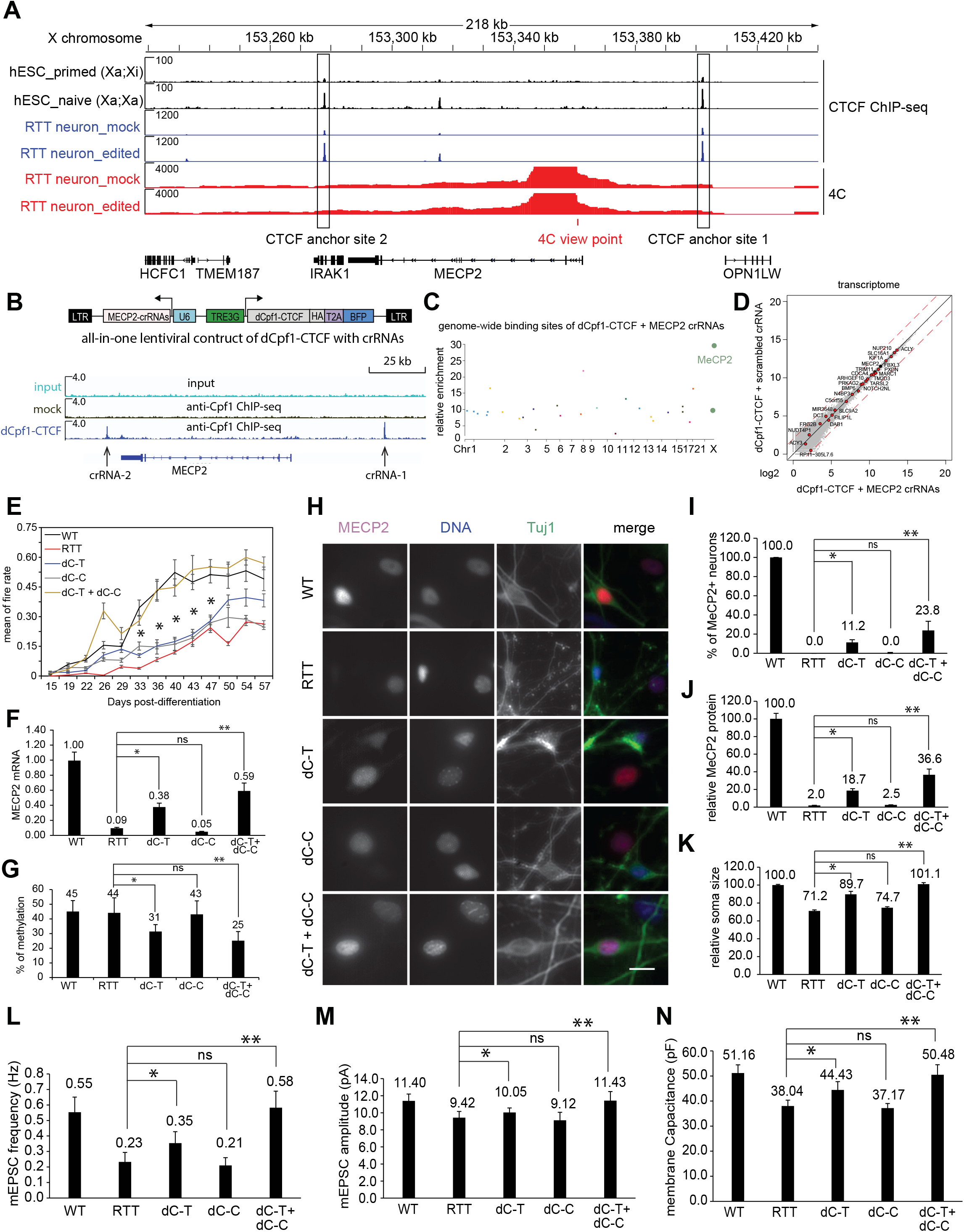
Multiplex epigenome editing of MECP2 to rescue RTT neurons. (**A**) Enrichment of CTCF binding around the *MECP2* locus in the neurons derived from mock (labeled RTT neuron_mock) or neuron derived from DNA methylation edited RTT hESCs (labeled as RTT neuron_edited), and primed or native RTT hESCs. 4C-seq was performed to reveal the genomic interactions of the *MECP2* promoter. The two CTCF anchor sites are labeled by rectangles in black. (**B**) Upper panel illustrates the all-in-one lentiviral construct to express Dox-inducible dCpf1-CTCF-HA and target crRNAs. Lower panel shows an anti-Cpf1 ChIP-seq using HEK293T cells transfected with empty vector or the dCpf1-CTCF construct with two crRNAs targeting the anchor sites in A. (**C**) A Manhattan plot showing 28 genome-wide binding sites of dCpf1-CTCF with the *MECP2* target crRNAs identified by anti-Cpf1 and anti-CTCF ChIP-seq. (**D**) Transcriptomes of cells transfected with dCpf1-CTCF + *MECP2* target crRNAs or dCpf1-CTCF + scrambled crRNA examined by RNA-seq. Red dots highlight the genes associated with the 28 dCpf1-CTCF binding sites identified in C. MECP2 is labeled with a black dot. The dashed red lines mark the 2-fold difference between the samples. (**E**) WT neurons, mock RTT neurons, or RTT neurons infected by lentiviral dCas9-Tet1/sgRNA-3 (labeled dC-T), or dCpf1-CTCF (dC-C), or both (dC-T + dC-C) were grown on the MEA plate for measurement of electrophysiological activities in a time course experiment. Shown is the mean ± SD of biological triplicates for each group of neurons. (**F**) *MECP2* mRNA quantity of the neurons in (E) was measured by qPCR on Day 57. Shown is the mean ± SD of biological triplicates for each group of neurons. (**G**) DNA methylation of the *MECP2* promoter in neurons in (E) was measured by Pyro-seq on Day 57. Shown is the average DNA methylation of CpGs within the targeted *MECP2* promoter region ± SD of three biological replicates. (**H**) Neurons in E were grown on mouse astrocytes to promote neuronal maturation and then IF stained with anti-MeCP2 and anti-Tuj1 antibodies. Scale bar: 50 um. (**I**) Percentages of neurons (Tuj1+) expressing MeCP2 protein within each group of samples described in H. (**J**) Quantification of MeCP2 protein amounts in the MeCP2-positive neurons in (H). (**K**) Quantification of soma sizes of the neurons in (H). Quantifications were done using ImageJ and are shown as the mean of relative percentages as compared to WT neurons, ± SD of two biological replicates. (**L to N**) The mEPSC frequency (**L**), mEPSC amplitude (**M**), and membrane capacitance (**N**) of neurons in (F). Shown is the mean ± SD of at least two biological repeats with more than 20 neurons for each condition. \**P* < 0.05, \*\**P* < 0.01, one-way ANOVA with Bonferroni correction.

To artificially recruit CTCF to these identified anchor sites, we generated a new epigenome manipulation tool by fusing a catalytically dead CRISPR/LbCpf1 (dCpf1) (*34, 35*) with CTCF to guide CTCF at specific genomic loci with target CRISPR RNA (crRNA). This all-in-one construct expresses dCpf1-CTCF driven by a Dox-inducible promoter and target crRNA array driven by a U6 promoter (**Figure 5B**, upper panel). Because Cpf1 recognizes TTTN as its protospacer adjacent motif (PAM) sequence and Cas9 recognizes NGG as PAM (*19, 34*), dCpf1-CTCF can be combined with dCas9-Tet1 to mediate multiplex epigenome editing in the same target cell without interfering each other. Using anti-Cpf1 ChIP-seq, we successfully targeted dCpf1-CTCF with crRNA-1 and crRNA-2 to the two identified CTCF anchor sites (**Figure 5B**, lower panel). Before applying this tool to RTT neurons, we examined its specificity and off-target effect. We first generated a control construct that expresses dCpf1-CTCF and a scrambled crRNA lacking a target sequence in the human genome. We performed anti-CTCF and anti-Cpf1 ChIP-seq using mock cells and cells expressing dCpf1-CTCF with either the *MECP2* target crRNAs or scrambled crRNA. We identified 28 binding sites for dCpf1-CTCF with *MECP2* target crRNAs by comparing the ChIP-seq peaks detected in these cells. Among these 28 binding sites, targeted MECP2 sites showed the highest enrichment (**Figure 5C** and **Data file S7**), suggesting an effective and specific targeting of dCpf1-CTCF. Next, we used RNA-seq to examine the expression of genes that were associated with these dCpf1-CTCF binding sites. None of these genes showed a significant change of expression (adjusted *P* value < 0.01, and fold change > 2; **Figure 5D** and **Data file S8-S9**), suggesting the off-target binding of dCpf1-CTCF did not result in transcriptional changes. Last, we compared the transcriptomes of cells expressing dCpf1-CTCF with *MECP2* target crRNAs or scrambled crRNA. Among 57,906 detected transcripts, three differentially expressed genes (adjusted *P* value < 0.01 and fold change > 2) were identified between these two groups (**Data file S10**). As dCpf1-CTCF did not bind to these three genes (**Figure 5C**), the expression changes of these three genes are not considered off-target effects by dCpf1-CTCF, but likely resulted from the transcriptional variation during cell culturing and passaging (*36*). In summary, the result from these ChIP-seq and RNA-seq experiments supports a specific targeting of dCpf1-CTCF to the *MECP2* sites without detectable off-target effects at the transcriptional level.

Next, we delivered dCas9-Tet1/sgRNA-3 alone (labeled dC-T), dCpf1-CTCF/crRNA alone (labeled dC-C), or both dC-T and dC-C into RTT neurons via lentiviral transduction as validated by qPCR analysis (**Figure S3)**. We then performed a multi-electrode array to examine the spontaneous electrophysiological activities of these infected neurons along the neuronal maturation process. RTT neurons expressing dC-T again showed partial rescue of the neuronal firing rate; neurons expressing dC-C behaved similarly as mock RTT neurons; however, neurons expressing both dC-T and dC-C displayed spontaneous neuronal activity indistinguishable from that of WT neurons (**Figure 5E)**, suggesting a complete rescue of neuronal activity in the multiplex edited RTT neurons. This greater degree of rescue on neuronal activity is due to more efficient reactivation of *MECP2* from Xi (59% of WT *MECP2* mRNA) as measured by qPCR (**Figure 5F)**, and more robust demethylation of the *MECP2* promoter (25% as the average methylation percentage of CpGs within the targeted *MECP2* promoter region) as measured by Pyro-seq (**Figure 5G)**. To evaluate the rescue effect on RTT-associated neuronal defects at a single neuron resolution, we cultured these neurons on mouse astrocytes to promote neuronal maturation and then performed a series of characterization experiments. Using the same quantification method as in Figure 4E-G (mouse astrocytes within each group as an internal control for MeCP2 protein abundance), we found that whereas 18.7% of MeCP2 protein was restored in 11.2% of RTT neurons after infection with dC-T alone, 36.6% of MeCP2 protein was detected in 23.8% of RTT neurons after infection with both dC-T and dC-C (**Figure 5H-6**), suggesting a higher restoration of MeCP2 at the protein level in multiplex-edited RTT neurons. Consistent with this result, the smaller soma size was completely rescued in these multiplex edited RTT neurons, but not by either dC-T or dC-C alone (**Figure 5K**). We also performed patch-clamp recording experiments using these neurons. The mEPSC frequency was increase from 0.23 Hz in mock RTT neuron to 0.58 Hz in multiplex edited RTT neurons similar to the 0.55 Hz in WT neurons (**Figure 5L**), and the mEPSC amplitude increased from 9.42 pA in mock RTT neurons to 11.43 pA in multiplex edited RTT neurons similar to the 11.40 in WT neurons (**Figure 5M**), suggesting an efficient restoration of excitatory synapses in these neurons. Last, the membrane capacitance was completely rescued in multiplex edited RTT neurons (**Figure 5N)**, consistent with the result of soma size analysis. These results demonstrate that a higher reactivation efficacy of MECP2 from Xi in RTT neurons can be achieved by multiplex epigenome editing that combines DNA methylation editing of the *MECP2* promoter by dCas9-Tet1 and insulation of the edited *MECP2* locus by targeted CTCF, resulting in better functional rescue of RTT-associated neuronal defects compared to DNA methylation editing alone in RTT neurons.

## Discussion

Our study demonstrates that DNA methylation editing of the *MECP2* promoter can efficiently reactivate the *MECP2* allele from Xi in RTT hESCs, and the cellular and electrophysiological defects of neurons derived from methylation edited RTT hESCs are functionally rescued. Previous screening studies identified several small chemical inhibitors targeting Aurora kinase A/B, Activin A Receptor Type 1, Janus kinase 2, phosphoinositide-dependent protein kinase-1, Polo-like kinase 2, RAD21 cohesin complex component, and ribonucleotide reductase that can slightly reactivate MECP2 from Xi with less than 1% of the active allele on Xa (*37-41*). Combination of ASO against XIST RNA with DNA methylation inhibitors such as 5-azacytidine or Decitabine resulted in a higher MECP2 reactivation with 2-5% of the active allele on Xa (*16*). However, this approach will likely cause the reactivation of other Xi genes in the treated cells. In contrast, the targeted reactivation approach by precise DNA methylation editing leads to a highly efficient reactivation (82% of the active allele at the protein level) in neurons derived from edited RTT hESCs and moderate reactivation (17.7% of the active allele at the protein level) in directly edited RTT neurons without affecting the expression of other genes on Xi or MECP2 on Xa.

DNA methylation editing alone by dCas9-Tet1 in RTT neurons resulted in a moderate reactivation of *MECP2* from Xi. This less efficient reactivation in neurons compared to hESCs is likely due to the lower demethylation efficacy achieved in directly edited neurons (33% methylation remained at the targeted MECP2 promoter) compared to 5% methylation in edited hESCs. We observed a similar trend when demethylation editing was performed in Fragile X syndrome iPSCs and iPSC-derived neurons (*24*). One reason for the different demethylation efficacies in RTT hESCs versus neurons is that both active (a DNA repair-based restoration into unmethylated cytosines) and passive demethylation (a DNA replication-dependent dilution of hemi-methylated cytosines) mechanisms can operate in dividing cells, whereas only active demethylation can be operated in post-mitotic neurons (*42*). In addition, other X chromosome inactivation epigenetic mechanisms such as XIST decoration might counteract DNA methylation editing to suppress MECP2 reactivation. These epigenetic suppressions are particularly effective in terminally differentiated and post-mitotic neurons to ensure X chromosome inactivation. Consistent with this hypothesis, combination of DNA methylation editing by dCas9-Tet1 with targeted CTCF insulation at the boundary of edited *MECP2* locus increased MECP2 reactivation from ∼30% to ∼59% of the active allele at the transcriptional level, resulting in a functional rescue of the RTT-associated neuronal defects.

In summary, our study provides a proof-of-concept to apply epigenome editing approach to rescue RTT neurons by reactivating the *MECP2* allele from Xi, suggesting a therapeutic strategy to treat RTT and potentially other X-linked human diseases. Limitations for this MECP2 reactivation-based strategy are 1) the current editors (dCas9-Tet1 and dCpf1-CTCF) are too large to fit into a single AAV vectors that are necessary for brain-wide gene delivery, 2) it is only applicable to female patients with RTT carrying a heterozygous loss-of-function mutants of MECP2, but not male patients with RTT lacking a wild type allele of MECP2, and 3) future study will be needed to test this epigenome editing strategy in animal models of RTT for rescue effect at behavioral levels.

## Supplementary materials

### Materials and Methods

Fig S1-S3

Tables S1-S3

Data files S1-S10

## Acknowledgments

We thank Michael Kissner at Columbia Stem Cell Initiative and Patti Wisniewski, Patrick Autissier, and Hanna Aharonov at Whitehead Institute for FACS. We thank Maisam Mitalipova and Dongdong Fu at Whitehead Institute for reagents and technical assistance. We thank Manu Ben-Johny at Columbia University for technical consultation on the electrophysiology study.

## Author contributions

XSL and RJ conceptualized the project. JQ, XG, BX, CX, JN, XT, and XSL contributed to methodology. JQ, XG, BX, CX, JN, XT, CHL, and XSL performed experiments. BX, XG, JN, XT, CHL, and XSL contributed to data visualization. XSL and RJ obtained funding. HC, XSL, and RJ supervised the project. XSL, RJ, JQ, XG, BX, CX, JN, XT, HC, and CHL wrote the manuscript.

## Data and materials availability

Cell lines, plasmids, and other materials are available upon request. Recipients of the cell lines or plasmids will need to execute a materials transfer agreement with Columbia University. The original NGS data associated with this study is available at the Gene Expression Omnibus (https://www.ncbi.nlm.nih.gov/geo/) under accession GSE214550.

## Materials and Methods

### Study design

The purpose of this study was to apply epigenome editing tools to specifically reactivate the wild type allele of *MECP2* from the inactive X chromosome and to examine whether the Rett syndrome-associated neuronal defects can be rescued in edited neurons. Based on our previous publications and power analysis results from pilot experiments, at least two biological replicates were used for each DNA methylation, transcription analysis, and a multi-electrode array assay using cultured hESCs or neurons. A sample size of more than 20 neurons was used for mEPSC measurements. Electrophysiological data were analyzed blind to genotype and treatment conditions. Morphological analyses were carried out with about 30 to 100 randomly selected neurons per treatment group. Image acquisition and analysis were carried out by different researchers in a double-blind manner, and automated analysis software pipelines were used to reduce human bias. This study was approved by Columbia University’s Human Embryonic and Human Pluripotent Stem Cell Research Committee.

### Statistical analysis

For all experiments described in the article, the experimentation, quantification, and analysis of data were performed with subjective unbiased methods, and the researchers conducting the experiments were blinded to genotype and treatment conditions. Data were tested for assumptions prior to use of parametric tests. For comparison of two groups, Student’s t tests were performed using Microsoft Excel software. For comparison of multiple groups, one-way analysis of variance (ANOVA) with Bonferroni post hoc correction was performed using GraphPad Prism software. Permutation test was used to compare mEPSC frequency results. Kolmogorov-Smirnov test was used to compare the cumulative distribution between two mEPSC amplitude datasets. Linear mixed modeling statistic test method was used to compare the neuronal morphological metrics.

Statistical parameters including the exact value of n, the definition of center, dispersion and precision measures (mean ± SD), and statistical significance are reported in the figure legends. Data was judged to be statistically significant when *P* < 0.05 or *P* < 0.01. *P* value was adjusted for False Discovery Rate in ChIP-seq, RNA-seq, and whole-genome bisulfite sequencing analyses.

## References and Notes

1. M. J. Lyst, A. Bird, Rett syndrome: a complex disorder with simple roots. Nat Rev Genet 16, 261–275 (2015); published online EpubMay (10.1038/nrg3897).

2. A. J. Sandweiss, V. L. Brandt, H. Y. Zoghbi, Advances in understanding of Rett syndrome and MECP2 duplication syndrome: prospects for future therapies. Lancet Neurol 19, 689–698 (2020); published online EpubAug (10.1016/S1474-4422(20)30217-9).

3. R. E. Amir, I. B. Van den Veyver, M. Wan, C. Q. Tran, U. Francke, H. Y. Zoghbi, Rett syndrome is caused by mutations in X-linked MECP2, encoding methyl-CpG-binding protein 2. Nat Genet 23, 185–188 (1999); published online EpubOct (10.1038/13810).

4. J. P. K. Ip, N. Mellios, M. Sur, Rett syndrome: insights into genetic, molecular and circuit mechanisms. Nat Rev Neurosci 19, 368–382 (2018); published online EpubJun (10.1038/s41583-018-0006-3).

5. W. Renthal, L. D. Boxer, S. Hrvatin, E. Li, A. Silberfeld, M. A. Nagy, E. C. Griffith, T. Vierbuchen, M. E. Greenberg, Characterization of human mosaic Rett syndrome brain tissue by single-nucleus RNA sequencing. Nat Neurosci 21, 1670–1679 (2018); published online EpubDec (10.1038/s41593-018-0270-6).

6. R. Chen, S. Akbarian, M. Tudor, R. Jaenisch, Deficiency of methyl-CpG binding protein-2 in CNS neurons results in a Rett-like phenotype in mice. Nat. Genet. 27, 327–331 (2001).

7. J. Guy, B. Hendrich, M. Holmes, J. E. Martin, A. Bird, A mouse Mecp2-null mutation causes neurological symptoms that mimic Rett syndrome. Nat. Genet. 27, 322–326 (2001); published online EpubMar (

8. N. Vashi, M. J. Justice, Treating Rett syndrome: from mouse models to human therapies. Mammalian genome : official journal of the International Mammalian Genome Society 30, 90–110 (2019); published online EpubJun (10.1007/s00335-019-09793-5).

9. E. S. Na, E. D. Nelson, E. T. Kavalali, L. M. Monteggia, The impact of MeCP2 loss-or gain-of-function on synaptic plasticity. Neuropsychopharmacology 38, 212–219 (2013); published online EpubJan (10.1038/npp.2012.116).

10. M. Shahbazian, J. Young, L. Yuva-Paylor, C. Spencer, B. Antalffy, J. Noebels, D. Armstrong, R. Paylor, H. Zoghbi, Mice with truncated MeCP2 recapitulate many Rett syndrome features and display hyperacetylation of histone H3. Neuron 35, 243–254 (2002); published online EpubJul 18 (10.1016/s0896-6273(02)00768-7).

11. E. Giacometti, S. Luikenhuis, C. Beard, R. Jaenisch, Partial rescue of MeCP2 deficiency by postnatal activation of MeCP2. Proc Natl Acad Sci U S A 104, 1931–1936 (2007); published online EpubFeb 6 (

12. J. Guy, J. Gan, J. Selfridge, S. Cobb, A. Bird, Reversal of neurological defects in a mouse model of Rett syndrome. Science 315, 1143–1147 (2007); published online EpubFeb 23 (

13. S. K. Garg, D. T. Lioy, H. Cheval, J. C. McGann, J. M. Bissonnette, M. J. Murtha, K. D. Foust, B. K. Kaspar, A. Bird, G. Mandel, Systemic delivery of MeCP2 rescues behavioral and cellular deficits in female mouse models of Rett syndrome. J Neurosci 33, 13612–13620 (2013); published online EpubAug 21 (10.1523/JNEUROSCI.1854-13.2013).

14. R. L. Adrianse, K. Smith, T. Gatbonton-Schwager, S. P. Sripathy, U. Lao, E. J. Foss, R. G. Boers, J. B. Boers, J. Gribnau, A. Bedalov, Perturbed maintenance of transcriptional repression on the inactive X-chromosome in the mouse brain after Xist deletion. Epigenetics Chromatin 11, 50 (2018); published online EpubAug 31 (10.1186/s13072-018-0219-8).

15. R. Tillotson, J. Selfridge, M. V. Koerner, K. K. E. Gadalla, J. Guy, D. De Sousa, R. D. Hector, S. R. Cobb, A. Bird, Radically truncated MeCP2 rescues Rett syndrome-like neurological defects. Nature 550, 398–401 (2017); published online EpubOct 19 (10.1038/nature24058).

16. N. B. Grimm, J. T. Lee, Selective Xi reactivation and alternative methods to restore MECP2 function in Rett syndrome. Trends Genet, (2022); published online EpubMar 2 (10.1016/j.tig.2022.01.007).

17. X. S. Liu, R. Jaenisch, Editing the Epigenome to Tackle Brain Disorders. Trends Neurosci 42, 861–870 (2019); published online EpubDec (10.1016/j.tins.2019.10.003).

18. R. Galupa, E. Heard, X-Chromosome Inactivation: A Crossroads Between Chromosome Architecture and Gene Regulation. Annu Rev Genet 52, 535–566 (2018); published online EpubNov 23 (10.1146/annurev-genet-120116-024611).

19. A. Pickar-Oliver, C. A. Gersbach, The next generation of CRISPR-Cas technologies and applications. Nat Rev Mol Cell Biol 20, 490–507 (2019); published online EpubAug (10.1038/s41580-019-0131-5).

20. X. S. Liu, H. Wu, X. Ji, Y. Stelzer, X. Wu, S. Czauderna, J. Shu, D. Dadon, R. A. Young, R. Jaenisch, Editing DNA Methylation in the Mammalian Genome. Cell 167, 233–247 e217 (2016); published online EpubSep 22 (10.1016/j.cell.2016.08.056).

21. A. C. Komor, A. H. Badran, D. R. Liu, CRISPR-Based Technologies for the Manipulation of Eukaryotic Genomes. Cell 169, 559 (2017); published online EpubApr 20 (10.1016/j.cell.2017.04.005).

22. J. A. Doudna, The promise and challenge of therapeutic genome editing. Nature 578, 229–236 (2020); published online EpubFeb (10.1038/s41586-020-1978-5).

23. A. L. Mattei, N. Bailly, A. Meissner, DNA methylation: a historical perspective. Trends Genet, (2022); published online EpubApr 30 (10.1016/j.tig.2022.03.010).

24. X. S. Liu, H. Wu, M. Krzisch, X. Wu, J. Graef, J. Muffat, D. Hnisz, C. H. Li, B. Yuan, C. Xu, Y. Li, D. Vershkov, A. Cacace, R. A. Young, R. Jaenisch, Rescue of Fragile X Syndrome Neurons by DNA Methylation Editing of the FMR1 Gene. Cell 172, 979–992 e976 (2018); published online EpubFeb 22 (10.1016/j.cell.2018.01.012).

25. T. W. Theunissen, M. Friedli, Y. He, E. Planet, R. C. O’Neil, S. Markoulaki, J. Pontis, H. Wang, A. Iouranova, M. Imbeault, J. Duc, M. A. Cohen, K. J. Wert, R. Castanon, Z. Zhang, Y. Huang, J. R. Nery, J. Drotar, T. Lungjangwa, D. Trono, J. R. Ecker, R. Jaenisch, Molecular Criteria for Defining the Naive Human Pluripotent State. Cell Stem Cell 19, 502–515 (2016); published online EpubOct 6 (10.1016/j.stem.2016.06.011).

26. R. Lister, M. Pelizzola, R. H. Dowen, R. D. Hawkins, G. Hon, J. Tonti-Filippini, J. R. Nery, L. Lee, Z. Ye, Q. M. Ngo, L. Edsall, J. Antosiewicz-Bourget, R. Stewart, V. Ruotti, A. H. Millar, J. A. Thomson, B. Ren, J. R. Ecker, Human DNA methylomes at base resolution show widespread epigenomic differences. Nature 462, 315–322 (2009); published online EpubNov 19 (10.1038/nature08514).

27. B. Tanasijevic, B. Dai, T. Ezashi, K. Livingston, R. M. Roberts, T. P. Rasmussen, Progressive accumulation of epigenetic heterogeneity during human ES cell culture. Epigenetics 4, 330–338 (2009); published online EpubJul 1 (10.4161/epi.4.5.9275).

28. S. Mekhoubad, C. Bock, A. S. de Boer, E. Kiskinis, A. Meissner, K. Eggan, Erosion of dosage compensation impacts human iPSC disease modeling. Cell Stem Cell 10, 595–609 (2012); published online EpubMay 4 (10.1016/j.stem.2012.02.014).

29. Y. Shi, P. Kirwan, J. Smith, H. P. Robinson, F. J. Livesey, Human cerebral cortex development from pluripotent stem cells to functional excitatory synapses. Nat Neurosci 15, 477-486, S471 (2012); published online EpubFeb 5 (10.1038/nn.3041).

30. Y. Shi, P. Kirwan, F. J. Livesey, Directed differentiation of human pluripotent stem cells to cerebral cortex neurons and neural networks. Nat Protoc 7, 1836–1846 (2012); published online EpubOct (10.1038/nprot.2012.116).

31. X. Tang, L. Zhou, A. M. Wagner, M. C. Marchetto, A. R. Muotri, F. H. Gage, G. Chen, Astroglial cells regulate the developmental timeline of human neurons differentiated from induced pluripotent stem cells. Stem Cell Res 11, 743–757 (2013); published online EpubSep (10.1016/j.scr.2013.05.002).

32. E. M. Darrow, M. H. Huntley, O. Dudchenko, E. K. Stamenova, N. C. Durand, Z. Sun, S. C. Huang, A. L. Sanborn, I. Machol, M. Shamim, A. P. Seberg, E. S. Lander, B. P. Chadwick, E. L. Aiden, Deletion of DXZ4 on the human inactive X chromosome alters higher-order genome architecture. Proc Natl Acad Sci U S A 113, E4504–4512 (2016); published online EpubAug 2 (10.1073/pnas.1609643113).

33. L. Giorgetti, B. R. Lajoie, A. C. Carter, M. Attia, Y. Zhan, J. Xu, C. J. Chen, N. Kaplan, H. Y. Chang, E. Heard, J. Dekker, Structural organization of the inactive X chromosome in the mouse. Nature 535, 575–579 (2016); published online EpubJul 28 (10.1038/nature18589).

34. B. Zetsche, J. S. Gootenberg, O. O. Abudayyeh, I. M. Slaymaker, K. S. Makarova, P. Essletzbichler, S. E. Volz, J. Joung, J. van der Oost, A. Regev, E. V. Koonin, F. Zhang, Cpf1 is a single RNA-guided endonuclease of a class 2 CRISPR-Cas system. Cell 163, 759–771 (2015); published online EpubOct 22 (10.1016/j.cell.2015.09.038).

35. B. Zetsche, M. Heidenreich, P. Mohanraju, I. Fedorova, J. Kneppers, E. M. DeGennaro, N. Winblad, S. R. Choudhury, O. O. Abudayyeh, J. S. Gootenberg, W. Y. Wu, D. A. Scott, K. Severinov, J. van der Oost, F. Zhang, Multiplex gene editing by CRISPR-Cpf1 using a single crRNA array. Nat Biotechnol 35, 31–34 (2017); published online EpubJan (10.1038/nbt.3737).

36. I. Carcamo-Orive, G. E. Hoffman, P. Cundiff, N. D. Beckmann, S. L. D’Souza, J. W. Knowles, A. Patel, D. Papatsenko, F. Abbasi, G. M. Reaven, S. Whalen, P. Lee, M. Shahbazi, M. Y. R. Henrion, K. Zhu, S. Wang, P. Roussos, E. E. Schadt, G. Pandey, R. Chang, T. Quertermous, I. Lemischka, Analysis of Transcriptional Variability in a Large Human iPSC Library Reveals Genetic and Non-genetic Determinants of Heterogeneity. Cell Stem Cell 20, 518–532 e519 (2017); published online EpubApr 6 (10.1016/j.stem.2016.11.005).

37. L. L. G. Carrette, C. Y. Wang, C. Wei, W. Press, W. Ma, R. J. Kelleher, 3rd, J. T. Lee, A mixed modality approach towards Xi reactivation for Rett syndrome and other X-linked disorders. Proc Natl Acad Sci U S A 115, E668–E675 (2018); published online EpubJan 23 (10.1073/pnas.1715124115).

38. S. Sripathy, V. Leko, R. L. Adrianse, T. Loe, E. J. Foss, E. Dalrymple, U. Lao, T. Gatbonton-Schwager, K. T. Carter, B. Payer, P. J. Paddison, W. M. Grady, J. T. Lee, M. S. Bartolomei, A. Bedalov, Screen for reactivation of MeCP2 on the inactive X chromosome identifies the BMP/TGF-beta superfamily as a regulator of XIST expression. Proc Natl Acad Sci U S A 114, 1619–1624 (2017); published online EpubFeb 14 (10.1073/pnas.1621356114).

39. D. Lessing, T. O. Dial, C. Wei, B. Payer, L. L. Carrette, B. Kesner, A. Szanto, A. Jadhav, D. J. Maloney, A. Simeonov, J. Theriault, T. Hasaka, A. Bedalov, M. S. Bartolomei, J. T. Lee, A high-throughput small molecule screen identifies synergism between DNA methylation and Aurora kinase pathways for X reactivation. Proc Natl Acad Sci U S A 113, 14366–14371 (2016); published online EpubDec 13 (10.1073/pnas.1617597113).

40. P. Przanowski, U. Wasko, Z. Zheng, J. Yu, R. Sherman, L. J. Zhu, M. J. McConnell, J. Tushir-Singh, M. R. Green, S. Bhatnagar, Pharmacological reactivation of inactive X-linked Mecp2 in cerebral cortical neurons of living mice. Proc Natl Acad Sci U S A 115, 7991–7996 (2018); published online EpubJul 31 (10.1073/pnas.1803792115).

41. H. M. Lee, M. B. Kuijer, N. Ruiz Blanes, E. P. Clark, M. Aita, L. Galiano Arjona, A. Kokot, N. Sciaky, J. M. Simon, S. Bhatnagar, B. D. Philpot, A. Cerase, A small-molecule screen reveals novel modulators of MeCP2 and X-chromosome inactivation maintenance. J Neurodev Disord 12, 29 (2020); published online EpubNov 10 (10.1186/s11689-020-09332-3).

42. H. Wu, Y. Zhang, Reversing DNA methylation: mechanisms, genomics, and biological functions. Cell 156, 45–68 (2014); published online EpubJan 16 (10.1016/j.cell.2013.12.019).

43. J. M. Dowen, Z. P. Fan, D. Hnisz, G. Ren, B. J. Abraham, L. N. Zhang, A. S. Weintraub, J. Schuijers, T. I. Lee, K. Zhao, R. A. Young, Control of cell identity genes occurs in insulated neighborhoods in mammalian chromosomes. Cell 159, 374–387 (2014); published online EpubOct 9 (10.1016/j.cell.2014.09.030).

44. X. Wu, D. A. Scott, A. J. Kriz, A. C. Chiu, P. D. Hsu, D. B. Dadon, A. W. Cheng, A. E. Trevino, S. Konermann, S. Chen, R. Jaenisch, F. Zhang, P. A. Sharp, Genome-wide binding of the CRISPR endonuclease Cas9 in mammalian cells. Nat Biotechnol 32, 670–676 (2014); published online EpubJul (10.1038/nbt.2889).

45. B. Langmead, S. L. Salzberg, Fast gapped-read alignment with Bowtie 2. Nat Methods 9, 357–359 (2012); published online EpubMar 4 (10.1038/nmeth.1923).

46. Y. Zhang, T. Liu, C. A. Meyer, J. Eeckhoute, D. S. Johnson, B. E. Bernstein, C. Nusbaum, R. M. Myers, M. Brown, W. Li, X. S. Liu, Model-based analysis of ChIP-Seq (MACS). Genome Biol 9, R137 (2008) 10.1186/gb-2008-9-9-r137).

47. S. Heinz, C. Benner, N. Spann, E. Bertolino, Y. C. Lin, P. Laslo, J. X. Cheng, C. Murre, H. Singh, C. K. Glass, Simple combinations of lineage-determining transcription factors prime cis-regulatory elements required for macrophage and B cell identities. Mol Cell 38, 576–589 (2010); published online EpubMay 28 (10.1016/j.molcel.2010.05.004).

48. F. Krueger, S. R. Andrews, Bismark: a flexible aligner and methylation caller for Bisulfite-Seq applications. Bioinformatics 27, 1571–1572 (2011); published online EpubJun 1 (10.1093/bioinformatics/btr167).

49. A. Akalin, M. Kormaksson, S. Li, F. E. Garrett-Bakelman, M. E. Figueroa, A. Melnick, C. E. Mason, methylKit: a comprehensive R package for the analysis of genome-wide DNA methylation profiles. Genome Biol 13, R87 (2012); published online EpubOct 3 (10.1186/gb-2012-13-10-r87).

50. A. S. Weintraub, C. H. Li, A. V. Zamudio, A. A. Sigova, N. M. Hannett, D. S. Day, B. J. Abraham, M. A. Cohen, B. Nabet, D. L. Buckley, Y. E. Guo, D. Hnisz, R. Jaenisch, J. E. Bradner, N. S. Gray, R. A. Young, YY1 Is a Structural Regulator of Enhancer-Promoter Loops. Cell 171, 1573–1588 e1528 (2017); published online EpubDec 14 (10.1016/j.cell.2017.11.008).

51. D. Kim, J. M. Paggi, C. Park, C. Bennett, S. L. Salzberg, Graph-based genome alignment and genotyping with HISAT2 and HISAT-genotype. Nat Biotechnol 37, 907–915 (2019); published online EpubAug (10.1038/s41587-019-0201-4).

52. Y. Liao, G. K. Smyth, W. Shi, featureCounts: an efficient general purpose program for assigning sequence reads to genomic features. Bioinformatics 30, 923–930 (2014); published online EpubApr 1 (10.1093/bioinformatics/btt656).

53. M. I. Love, W. Huber, S. Anders, Moderated estimation of fold change and dispersion for RNA-seq data with DESeq2. Genome Biol 15, 550 (2014)10.1186/s13059-014-0550-8).

