## Supplemental Material for "Multiplex Epigenome Editing of *MECP2* to Rescue Rett Syndrome Neurons"

### Materials and Methods

#### Plasmid design and construction

PCR amplified TRE promoter from 138-dCas9-Dnmt3a (Addgene plasmid: 84570), tagBFP (synthesized gene block from IDT), and U6-crRNA array (synthesized gene block from IDT) were cloned into FUW vector (Addgene plasmid: 14882) via Gibson Assembly (NEB) as a modified FUW lentiviral vector construct. PCR amplified CTCF from pDONR223_CTCF_WT (Addgene plasmid: 81789) and human codon optimized catalytically inactive LbCpf1 (dLbCpf1) with mutation of D832A (synthesized gene blocks from IDT) were cloned into modified FUW vector construct via Gibson Assembly to package lentiviruses. The sgRNA expression plasmids were cloned by inserting annealed oligos into modified pgRNA plasmid (Addgene plasmid: 44248) with AarI site. All constructs were sanger sequenced for validation before transfection. Primer information for sgRNA design and construction is listed in Table S1. Related plasmids have been deposited into Addgene plasmid database for request (Addgene ID: 194885-194903).

#### Cell culture and lentivirus production

Human ESCs were cultured either with mTeSR1 medium (STEMCELL, #85850) or on irradiated mouse embryonic fibroblasts (MEFs) with standard hESCs medium: [DMEM/F12 (Invitrogen) supplemented with 15% fetal bovine serum (Gibco HI FBS, 10082-147), 5% KnockOut Serum Replacement (Invitrogen), 2 mM L-glutamine (MPBio), 1% nonessential amino acids (Invitrogen), 1% penicillin-streptomycin (Lonza), 0.1 mM β-mercaptoethanol (Sigma) and 4 ng/ml FGF2 (R&D systems)]. Lentiviruses expressing dCas9-Tet1-P2A-BFP, dCas9-dTet1-P2A-BFP, dCpf1-CTCF-P2A-BFP, rtTA and sgRNAs were produced by transfecting HEK293T cells with FUW constructs or sgRNA constructs together with standard packaging vectors (pCMV-dR8.74 and pCMV-VSVG) followed by ultra-centrifugation-based concentration. Virus titer (T) was calculated based on the infection efficiency for HEK293T cells, where $T = (P*N) / (V)$, T = titer (TU/ul), P = % of infection positive cells according to the fluorescence marker, N = number of cells at the time of transduction, V = total volume of virus used. Note TU stands for transduction unit.

#### Immunohistochemistry, microscopy, and image analysis

Human ESCs and neurons were fixed with 4% paraformaldehyde (PFA) for 10 min at room temperature. Cells were permeabilized with PBST (1 x PBS solution with 0.1% Triton X-100) before blocking with 10% Normal Donkey Serum (NDS) in PBST. Cells were then incubated with appropriately diluted primary antibodies in PBST with 5% NDS for 1 hours at room temperature or 12 hours at 4 °C, washed with PBST for 3 times at room temperature and then incubated with desired secondary antibodies in TBST with 5% NDS and DAPI to counter stain the nuclei. Cells were washed 3 times with PBST before mounted onto slides with Fluoromount G (SouthernBiotech). Sections were washed 3 times with PBST before slide mounting. The following antibodies were used in this study: Chicken anti-GFP (1:1000, Aves Labs), Mouse anti-Cas9 (7A9, 1:1000, Active Motif), Chicken anti-MAP2 (1:1000, Encor Biotech), and Mouse anti-Tuj1 (1:1000, Biolegend). Images were captured on a Nikon Eclipse TS2R microscope and processed with NIS-Elements software, ImageJ/Fiji, and Adobe Photoshop. For imaging-based quantification, unless otherwise specified, 3-5 representative images were quantified and data were plotted as mean ± SD with Excel or Graphpad.

**FACS analysis**

To isolate the infection-positive cells after lentiviral transduction, the treated cells were dissociated with Accutase and single-cell suspensions were prepared in growth medium subject to a BD FACSAria cell sorter according to the manufacture's protocol at the Whitehead Institute Flow Cytometry Core and Columbia Stem Cell Initiative Flow Cytometry Core. Data were analyzed with FlowJo software.

**Western blot**

Cells were lysed by RIPA buffer with proteinase inhibitor (Invitrogen), and subject to standard immunoblotting analysis. Mouse anti-Cas9 (1:1000, Active Motif), mouse α-Tubulin (1:10,000, Sigma), rabbit anti-MeCP2 (1:1000, Diagenode) antibodies were used.

**RT-qPCR**

Cells were harvested using Trizol followed by Direct-zol (Zymo Research), according to manufacturer's instructions. RNA was converted to cDNA using First-strand cDNA synthesis (Invitrogen SuperScript III). Quantitative PCR reactions were prepared with SYBR Green (Invitrogen), and performed in Applied Biosystems QuantStudio Real-Time PCR System. Primer information for RT-qPCR is listed in Table S2.

**Multi-electrode array recording**

Two- or four-week-old differentiating neuronal cultures were dissociated using Accutase and 5 X $10^5$ cells were plated on each single well of the PEI-coated Axion Biosystems plate (#M768-GL1-30Pt200) or MaxWell Two 6-well plate (#MX2-S-6W). Recordings of spontaneous activities during a 5-minute period were performed on days indicated. Biological triplicates for each type of neurons were included.

**ChIP assay**

ChIP experiment was performed as previously described (*43*). Briefly, cells were cross-linked by 1% formaldehyde in the medium for 10 min in room temperature, and then quenched by adding 0.125 M Glycine for 5 min. Collected cells were washed with PBS twice, and then re-suspended in 3.5 ml of sonication buffer. Sonication was performed for 10 cycles with 0.5 min pulse on and 1 min rest, and 24 watts in ice-water mixture. Then cell lysate was spun down with 14,000 x rpm for 10 min at 4 °C. 50 ul of supernatant was saved as input for gDNA. 10 ul of anti-Cas9 antibody (Active Motif) was added and incubate overnight at 4 °C. 50 ul protein G dynabeads was added into antibody-cell lysate mixture and incubate overnight at 4 °C. Then beads were washed with sonication buffer with high salt (500 mM NaCl), LiCl wash buffer, and TE buffer. Bound protein-DNA complex was eluted from beads by incubation in a 65 °C oven for 15 min, and then reverse cross-linked under 65 °C over-night. The bound DNA was purified with Qiagen QIAquick PCR Purification Kit, and then subject to qPCR analysis or sequencing.

**ChIP-seq peak calling method**

Sequencing data was analyzed with a previously reported method (*44*) with modifications. In brief, Trim Galore (https://www.bioinformatics.babraham.ac.uk/projects/trim_galore/) was used to remove adaptor and low-quality reads. The high-quality reads were mapped to human GRCh38 version genome using Bowtie2 with default parameters (*45*). To match sequencing depth, 7 million reads were randomly sampled from each library. Peaks were called by MACS2 (*46*). For dCas9-Tet1 with sgRNA-3 sample, input and dCas9-Tet1 with scrambled sgRNA were used as control separately to call peaks. Then the unique peaks were defined as binding sites for dCas9-Tet1 with sgRNA-3. Off-target effects were analyzed with a previously reported method (*44*). Peaks were annotated to its nearest genes by Homer (*47*) using ensemble GRCh38 version genome sequence and GTF file. For dCpf1-CTCF with MECP2 crRNAs sample, high-quality reads were mapped to human hg19 version genome using Bowtie2 with default parameters. Duplicated reads were removed by Picard (https://broadinstitute.github.io/picard/). MACS2 was used for peak calling. For each sample, its own input was used as a control, and q value (adjusted *P* value for the False Discovery Rate) for peak detection was set as $10^{-10}$. After removed the overlapped peaks with its own mock control, the overlapping peaks between anti-CTCF and anti-Cpf1 ChIP-seq were defined as binding sites for dCpf1-CTCF with MECP2 crRNAs. Reads mapped to the plasmid of dCpf1-CTCF in the transfected cells were manually removed.

**ChIP-BS-seq**

Anti-Cas9 ChIP experiment was performed as described above. The BS conversion and sequencing library preparation were performed according to the instructions by Pico Methyl-Seq Library Prep Kit (Zymo Research, D5455).

**ChIP-BS-seq analysis method**

Adaptor sequences and low-quality reads were removed by Trim Galore. The remaining high-quality reads were mapped to human GRCh38 version genome by Bismark (*48*) with the parameters "--non_directional --un --ambiguous --bowtie2 -N 1 -p 2 -score_min L, -6, -0.3." The methylation level in overlapped peak regions were called by methylpy (*48*). CpG sites with at least 5 reads coverage were used to calculate the DNA methylation level in dCas9-Tet1/sgRNA-3 and dCas9-dTet1/sgRNA-3 samples. For the regions without enough read coverage, pyro-sequencing was used to detect the DNA methylation percentages in both samples.

**Whole genome bisulfite sequencing and analysis method**

29-R hESCs expressing dCas9-Tet1 or dCas9-dead Tet1 with *MECP2* target sgRNA-3 were used to extract genomic DNA and subject to sequencing library preparation according to the instructions by Pico Methyl-Seq Library Prep Kit (Zymo Research, D5455). Three biological replicates were included for each group. The sequencing was performed by Novogene. Adaptor sequences and low-quality reads were removed by Trim Galore. The remaining high-quality reads were mapped to human GRCh38 version genome by Bismark with the parameters "--non_directional --un --ambiguous --bowtie2 -N 1 -p 2 -score_min L, -6, -0.3." The differentially methylated CpG (DMC) was called by methylkit (*49*). The cutoff for DMCs was as following: each CpG site had at least 5 reads coverage and only the CpG sites were covered in at least two replications were used to calculate the DNA methylation percentages in dCas9-Tet1/sgRNA-3 and dCas9-deadTet1/sgRNA-3 samples, q value (adjusted *P* value for the False Discovery Rate) smaller than 0.01, DNA methylation percentage change larger than 20%. Chiseq test was used for the methylation differences calculation. Based on the DMCs, differentially methylated regions (DMR) were called based on following cutoff: at least includes 3 DMCs with the same DNA methylation change direction, and the distance between each DMCs is less than 250bp. The genes annotated in the differentially methylated regions were defined as differentially methylated genes (DMG).

**4C-seq and analysis**

4C-seq experiment was performed as described in (*50*). Briefly, neuronal cultures were dissociates using Accutase and 5 million cells were resuspended in 5 mL 10% FBS in PBS. 5 mL of 4% formaldehyde in 10% FBS in PBS was added and cells were crosslinked for 10 minutes. Glycine was added to a final concentration of 0.125M and cells were centrifuged at 300 rcf for 5 minutes. Cells were washed twice with PBS, then snap frozen and stored at -80 C.

Pellets were gently resuspended in Hi-C lysis buffer (10 mM Tris-HCl pH 8, 10 mM NaCl, 0.2% Igepal CA-630) with 1x cOmplete protease inhibitors (Roche 11697498001). Cells were incubated on ice for 30 minutes then washed once with 500 mL of ice-cold Hi-C lysis buffer. Pellets were resuspended in 50 μL of 0.5% SDS and incubated at 62 C for 7 minutes. 145 μL of H2O and 25 μL of 10% Triton X-100 were added and tubes incubated at 37 C for 15 minutes. 25 μL of the appropriate 10X New England Biolabs restriction enzyme buffer and 200 units of DpnII (NEB, R0543M) were added and the chromatin was incubated at 37 C in a thermomixer at 500 rpm for four hours, 200 additional units of DpnII was added and the reaction was incubated overnight at 37 C in a thermomixer at 500 rpm, then a further 200 units of DpnII was added and the reaction was incubated another four hours at 37 C in a thermomixer at 500 rpm. Restriction enzyme was inactivated by heating to 65 C for 20 minutes while shaking at 500 rpm. Proximity ligation was performed in a total of 1200 μL with 2000 units of T4 DNA ligase (NEB M020) for six hours at room temperature with rotation. After ligation samples were spun down for 5 minutes at 2500 rcf and resuspended in 300 μL 10mM Tris-HCl, 1% SDS and 0.5 mM NaCl with 1000 units of Proteinase K. Cross-links were reversed by incubation overnight at 65 C. Samples were then phenol-chloroform extracted and ethanol precipitated, and the second digestion was performed overnight with 50 units of CviQI (NEB R0639S) in 450 μL of 1X NEBuffer. Samples were phenol-chloroform extracted and ethanol precipitated, and the second ligation was performed in 14 μL total with 6700 units of T4 DNA ligase (NEB M020) at 16 C overnight with rotation. Samples were ethanol precipitated, resuspended in 500 μL of 10 mM Tris-HCl pH 7.5, and purified with a QIAGEN PCR purification kit. Purified 4C template DNA was resuspended at 50 ng/L in 10 mM Tris-HCl pH 7.5.

To amplify 4C libraries, PCR amplification was performed with 16 x 50 μL PCR reactions using Roche Expand Long Template polymerase (Roche 11759060001). Reaction conditions are as follows: 11.2 μL Roche Expand Long Template Polymerase, 80 μL of 10 X Roche Buffer 1, 16 μL of 10 mM dNTPs (Promega PAU1515), 112 μL of 10 M forward primer, 112 μL of 10 M reverse primer, 200 ng 4C template DNA, and H2O until 800 μL total. Reactions were mixed and then distributed into 16 x 50 μL reactions for amplification. Cycling conditions were: 2 minutes at 94 C, 10 seconds at 94 C, 1 minute at 63 C, 30 seconds at 68 C, repeat steps 2-4 but decrease annealing temperature by one degree, until 53 C is reached at which point the reaction is cycled an additional 15 times at 53 C, after 25 total cycles are performed the reaction is held for 5 minutes at 68 C and then 4 C. Libraries were cleaned-up using a Roche High Pure PCR purification kit (Roche 11732676001) using 4 columns per library. Purified 4C-seq libraries were resuspended in 120 μL of 10 mM Tris-HCl pH 7.5. Purified 4C-seq libraries were quantified and sequenced on using an Illumina HiSeq 2500 instrument using 40 bp single-end reads.

**4C-seq primers**

DC26955686-forward

AATGATACGGCGACCACCGAACACTCTTTCCCTACACGACGCTCTTCCGATCT[nnn]TGTCT
CTGCTGTGCGGGATC

[nnn] is 2 or 3 base multiplexing barcode to distinguish samples

DC26955686-reverse

CAAGCAGAAGACGGCATACGAtcatgaccaaactgaacaga

DC26955706-foward

AATGATACGGCGACCACCGAACACTCTTTCCCTACACGACGCTCTTCCGATCT[nnn]CTACA
CCTGCAACATAGATC

[nnn] is 2 or 3 base multiplexing barcode to distinguish samples

DC26955706-reverse

CAAGCAGAAGACGGCATACGAGATGTAGGGCATAGAGCAAG

**4C-seq Data Analysis**

4C-seq samples were processed using fourfold (https://github.com/younglab/fourfold). Samples were first processed by removing their associated read primer sequences from the 50 end of each FASTQ read. To improve mapping efficiency of the trimmed reads by making the read longer, the restriction enzyme digest site was kept on the trimmed read. After trimming the reads, the reads were mapped using bowtie with options –k 1 –m 1 against the hg19 genome assembly. All unmapped or repetitively mapping reads were discarded from further analysis. The hg19 genome was then ‘‘digested’’ in silico according to the restriction enzyme pair used for that sample to identify all the fragments that could be generated by a 4C experiment given a restriction enzyme pair. All mapped reads were assigned to their corresponding fragment based on where they mapped to the genome.

**Bisulfite conversion and Pyro-sequencing**

Bisulfite conversion of DNA was established using the EpiTect Bisulfite Kit (Qiagen) following the manufacturer's instructions. Pyro-seq of all bisulfite converted genomic DNA samples were performed with PyroMark Q48 Autoprep (Qiagen) according to the manufacturer's instructions. Primer information for pyro-seq is listed in Table S3.

**RNA-seq analysis**

Adaptor sequences and low-quality reads were trimmed by Trim Galore. Using hisat2 (*51*), the high-quality reads were mapped to human GRCh38 version genome, tdTomato and GFP sequence. The read counts were obtained using featureCounts (*52*). Reads were normalized with DESeq2 (*53*). To avoid $\log_2 0$ as undefined value, the average number of reads for each gene was increased by one.

**Patch-clamp recording and data analysis**

Functionally mature stem cell-derived human neuron cultures were generated through seeding neural progenitor cells or neurons on a supporting layer of mouse astrocytes (*31*). After 8 weeks for neural progenitor cells or 4 weeks for neurons in culture, coverslips containing human neurons were transferred to a recording chamber and perfused with bath solution that consists of (in mM): 120 NaCl, 30 glucose, 25 HEPES, 5 KCl, 2 $CaCl_2$, 1 $MgCl_2$, pH adjusted to 7.3 with NaOH, and osmolality adjusted to 320. To record mini excitatory post-synaptic currents (mEPSC), we used potassium gluconate internal pipette solution consists of (in mM): 125 KGluconate, 10 KCl, 5 EGTA, 10 HEPES, 10 Tris-phosphocreatine, 4 MgATP, and 0.5 $Na_2GTP$; pH was adjusted to 7.3 with KOH, and osmolality adjusted to 305. After a whole-cell patch clamp recording configuration is established, the membrane potential was held at -70 mV, and 0.5 μM of TTX and 50 μM Picrotoxin were applied through bath solution perfusion to block sodium channels and $GABA_A$ receptors for recording mEPSC. The membrane capacitance of recorded neurons was estimated with the membrane test function of the Clampex software. Electrophysiological signals were amplified with Multiclamp 700A patch-clamp amplifier and data were collected with Clamp 9 software (Molecular Devices), sampled at 10 kHz and filtered at 1 kHz. Off-line data analyses of mEPSC synaptic events were performed with MiniAnalysis software (Synaptosoft). All experiments were performed at room temperature.

**Supplementary Figures**

### Figure S1

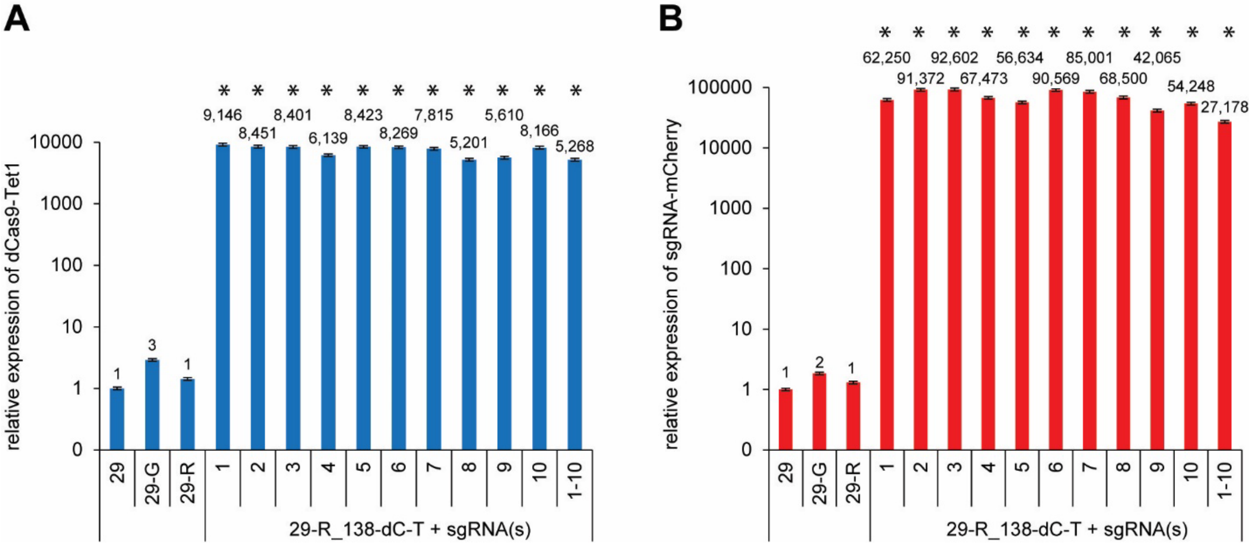

**Figure S1**. **Expression of dCas9-Tet1 and sgRNAs in edited 29-R hESCs**. A Dox-inducible dCas9-Tet1 expression 29-R hESCs were infected with lentiviruses expressing individual sgRNA or the mixture of 10 sgRNAs with mCherry as a fluorescent marker targeting the *MECP2* promoter region that overlaps with the differentially methylated region (DMR) between female and male hESCs. The expression of dCas9-Tet1 was examined in (**A**) and the expression of sgRNA-mCherry in (**B**) by qPCR. Shown is the mean percentage ± SD of three biological replicates. *$P$ < 0.05, one-way analysis of variance (ANOVA) with Bonferroni correction.

### Figure S2

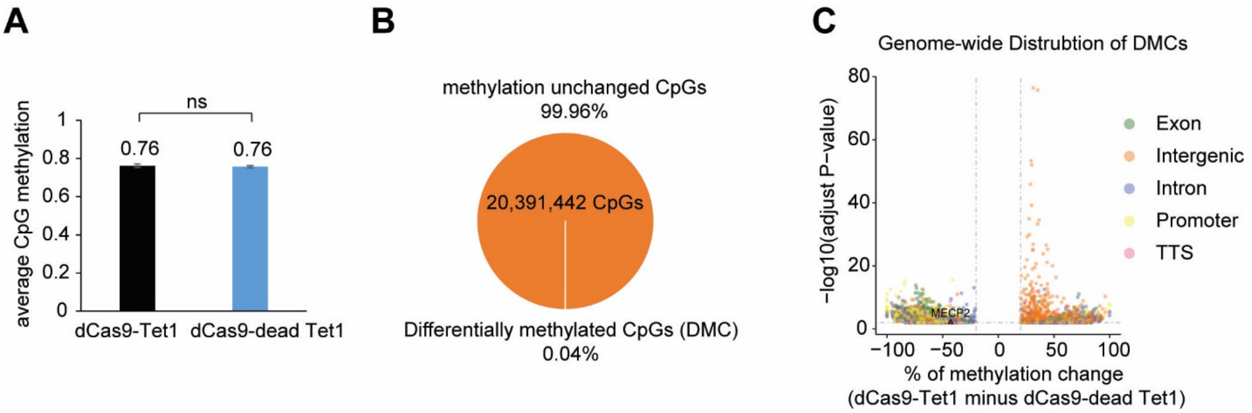

**Figure S2**. **Whole-genome bisulfite sequencing of 29-R cells expressing dCas9-Tet1 or dCas9-dead Tet1 with sgRNA-3**. A Dox-inducible dCas9-Tet1 or dCas9-dead Tet1 expression 29-R hESCs were infected with lentiviruses expressing *MECP2* target sgRNA-3. After treatment with Dox, the genomic DNA were extracted and subject to bisulfite conversion and then prepared as a sequencing library. (**A**) The averages of cytosine methylation in the CpG context for these two groups of cells. ns means not significant (*P* value > 0.05). (**B**) The percentages of methylation unchanged CpGs and differentially methylated CpGs (DMC) between these two groups of cells. adjusted *P* value < 0.01 and change of methylation > 20% for DMCs. (**C**) The genomewide distribution of DMCs. Three biological replicates for each group.

### Figure S3

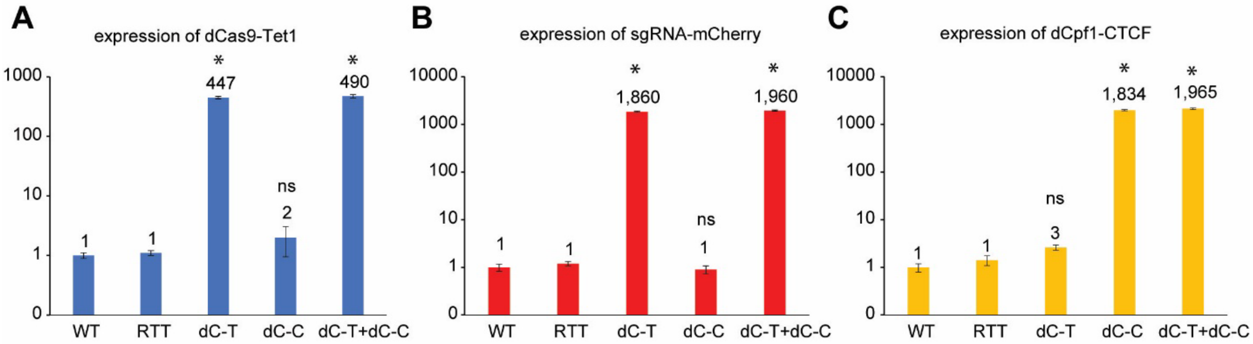

**Figure S3**. **Expression of dCas9-Tet1, sgRNAs, and dCpf1-CTCF in edited RTT cells**. Gene expression analysis of WT, RTT, or RTT neurons expressing dCas9-Tet1/sgRNA-3 (labeled as dC-T), or dCpf1-CTCF (labeled as dC-C), or both (labeled as dC-T + dC-C). Shown is the mean ± SD of biological replicates for each type of neurons. The expression of dCas9-Tet1 was examined in (**A**), the expression of sgRNA-mCherry in (**B**), and the expression of dCpf1-CTCF in (**C**). *$P < 0.05$, one-way ANOVA with Bonferroni correction.

Supplemental Tables

**Table S1. Target sgRNA and crRNA sequences.**

| **sgRNAs targeting the MECP2 promoter** | **5' to 3'** |
|---|---|
| SL-586_hMeCP2_DMR_ sgRNA-1_For | TTGG AGCAGCAAAGTTGCCCACCC |
| SL-587_hMeCP2_DMR _sgRNA-1_Rev | AAAC GGGTGGGCAACTTTGCTGCT |
| SL-588_hMeCP2_DMR_ sgRNA-2_For | TTGG TAGTGATATTGAGAAAATGT |
| SL-589_hMeCP2_DMR _sgRNA-2_Rev | AAAC ACATTTTCTCAATATCACTA |
| SL-590_hMeCP2_DMR_ sgRNA-3_For | TTGG CAGCCAATCAACAGCTGGAG |
| SL-591_hMeCP2_DMR _sgRNA-3_Rev | AAAC CTCCAGCTGTTGATTGGCTG |
| SL-592_hMeCP2_DMR_ sgRNA-4_For | TTGG GCCATCACAGCCAATGAC |
| SL-593_hMeCP2_DMR _sgRNA-4_Rev | AAAC GTCATTGGCTGTGATGGC |
| SL-594_hMeCP2_DMR_ sgRNA-5_For | TTGG AGGAGGAGAGACTGTGAGT |
| SL-595_hMeCP2_DMR _sgRNA-5_Rev | AAAC ACTCACAGTCTCTCCTCCT |
| SL-596_hMeCP2_DMR_ sgRNA-6_For | TTGG GGAGGGGGAGGGTAGAGAGG |
| SL-597_hMeCP2_DMR _sgRNA-6_Rev | AAAC CCTCTCTACCCTCCCCCTCC |
| SL-598_hMeCP2_DMR_ sgRNA-7_For | TTGG GGGAGGAAGAGGGGCGTC |
| SL-599_hMeCP2_DMR _sgRNA-7_Rev | AAAC GACGCCCCTCTTCCTCCC |
| SL-600_hMeCP2_DMR_ sgRNA-8_For | TTGG TGAGAGCTCAGGAGCCCTTG |
| SL-601_hMeCP2_DMR _sgRNA-8_Rev | AAAC CAAGGGCTCCTGAGCTCTCA |
| SL-602_hMeCP2_DMR_ sgRNA-9_For | TTGG CCTACTTGTTCCTGCTAGAT |
| SL-603_hMeCP2_DMR _sgRNA-9_Rev | AAAC ATCTAGCAGGAACAAGTAGG |
| SL-604_hMeCP2_DMR_ sgRNA-10_For | TTGG AGGTGGTTATAGTTCCCATC |
| SL-605_hMeCP2_DMR _sgRNA-10_Rev | AAAC GATGGGAACTATAACCACCT |
| **crRNAs targeting the CTCF anchor sites** | |
| crRNA for CTCF anchor site 1 | aggacatggcccaggcccgctca |
| crRNA for CTCF anchor site 2 | cttctctctatgtgaagggggtc |

**Table S2. qPCR primers.**

| | |
|---|---|
| SL-17_hMECP2 qPCR targeting Exon 4_For | GCAGAAAAGTACAAACACCGAGGG |
| SL-18_hMECP2 qPCR targeting Exon 4_Rev | GACAACAGCTGCCTTTATTCTTGTTGG |
| SL-481_eGFP_qPCR_2_For | AGAACGGCATCAAGGTGAAC |
| SL-482_eGFP_qPCR_2_Rev | TGCTCAGGTAGTGGTTGTCG |
| SL-838_tdTomato_qPCR_For | CACCATCGTGGAACAGTACG |
| SL-837_tdTomato_qPCR_Rev | atgacggccatgttgttgt |
| SL-295_dCas9_PCR screen_For | CCTTCGAGAAGAACCCAATTGAC |
| SL-356_dCas9_PCR screen_Rev | GGCTTGCAAGGTAGAGGAAATTC |
| SL-846_LbCpf1_qPCR_1_For | ctggtggaggacgagaagag |
| SL-847_LbCpf1_qPCR_1_Rev | ccggaacaggctgatgtaat |
| SL-711_mCherry_qPCR_For | cactacgacgctgaggtcaa |
| SL-712_mCherry_qPCR_Rev | gtgggaggtgatgtccaact |
| SL-899_hMap2_qPCR_For | TTGGTGCCGAGTGAGAAGAA |
| SL-900_hMap2_qPCR_Rev | GGTCTGGCAGTGGTTGGTTAA |
| SL-857_hHPRT_qPCR_1_For | CCTGGCGTCGTGATTAGTGAT |
| SL-858_hHRPT_qPCR_1_Rev | AGACGTTCAGTCCTGTCCATAA |
| SL-859_hPRPS1_qPCR_1_For | ATCTTCTCCGGTCCTGCTATT |
| SL-860_hPRPS1_qPCR_1_Rev | TGGTGACTACTACTGCCTCAAA |
| SL-861_hCOL4A5_qPCR_1_For | TGGACAGGATGGATTGCCAG |
| SL-862_hCOL4A5_qPCR_1_Rev | GGGGACCTCTTTCACCCTTAAAA |
| SL-863_hCASK_qPCR_1_For | TTGAAATCGTAAAGCGAGCTGA |
| SL-864_hCASK_qPCR_1_Rev | CAGTAGCGTAGAGCTTCCAGTA |
| SL-865_hRPGR_qPCR_1_For | TTCCGAAGGGCAAATTGGTTT |
| SL-866_hRPGR_qPCR_1_Rev | ACTTCCCATTCTCAGGTTCTCC |
| SL-867_hPDHA1_qPCR_1_For | TGGTAGCATCCCGTAATTTTGC |
| SL-868_hPDHA1_qPCR_1_Rev | ATTCGGCGTACAGTCTGCATC |
| SL-869_hHCCS_qPCR_1_For | TGATCCGGTTTGGAGGGAAAG |
| SL-870_hHCCS_qPCR_1_Rev | CAAAAGGCAACTCATACCCCAT |

**Table S3. Pyrosequencing primers.**

| | |
|---|---|
| **MECP2 promoter BS-seq** | |
| SL-813_hMECP2 promoter_area c_For | GAGGGGGAGGGTAGAGAG |
| SL-814_hMECP2 promoter_area c_Rev_Biotin | CTCCCTCCTCTCCAAAAAAAAACTATAATA |
| SL-815_hMECP2 promoter_area c_Seq | GGGAGGGTAGAGAGG |
| SL-816_hMECP2 promoter_area b_For | GGGTAGAGGGGGGTAGAAATT |
| SL-817_hMECP2 promoter_area b_Rev_Biotin | ACCCCCACCTCTCCCTAAAT |
| SL-818_hMECP2 promoter_area b_Seq | AGAGTTTAGGAGTTTTTGT |
| SL-819_hMECP2 promoter_area a_For | GAGTTGTGGGATTTAGAATATAATGT |
| SL-820_hMECP2 promoter_area a_Biotin | CTCCTTCTCCCCCATTCCATAAATTTC |
| SL-821_hMECP2 promoter_area a_Seq | GTTAGATGGGGAAAGG |
| SL-843_hMECP2 promoter_area d_For_Biotin | AGTTTTTTTTTAGAGAGGAGGGAG |
| SL-844_hMECP2 promoter_area d_Rev | TCCCAACCCCACCATCACAAC |
| SL-845_hMECP2 promoter_area d_Seq | ACCATCACAACCAAT |

The following Excel data files are available online:

**Data file S1**. Genome-wide binding sites of dCas9-Tet1 with sgRNA-3.

**Data file S2**. DNA methylation of dCas9-Tet1 binding sites.

**Data file S3**. RNA-seq of dCas9-Tet1 and dCas9-dead Tet1 samples

**Data file S4**. Expression of genes associated with dCas9-Tet1 binding sites

**Data file S5**. DEGs between dCas9-Tet1 and dCas9-dead Tet1 samples

**Data file S6**. Expression of genes associated with DMCs

**Data file S7**. Genome-wide binding sites of dCpf1-CTCF with MECP2 crRNAs

**Data file S8**. RNA-seq of dCpf1-CTCF with MECP2 crRNA or Scrambled crRNA

**Data file S9**. Expression of genes associated with dCpf1-CTCF binding sites

**Data file S10**. DEGs between dCpf1-CTCF with MECP2 crRNAs and Scrambled crRNA
